# Combination of enhanced gene activation and cell death causes lethal inflammatory pulmonary fibrosis

**DOI:** 10.64898/2026.09.03.749194

**Authors:** Julia Saggau, Christine Kiefer, Debora Bonasera, Luisa Schmidt, Louisa Grauvogel, Paulina Engel, Santiago Serrano-Sáenz, Craig Thrussell, Hassan Rakhsh-Khorshid, Konstantinos Kelepouras, Zsolt Megyesfalvi, Balazs Dome, Marcus Krüger, Antonella Montinaro, Nima Abedpour, Eva Rieser, Gianmaria Liccardi, Henning Walczak

## Abstract

The current paradigm of inflammatory disease etiology is that it is caused by an aberrant increase in either gene activation or cell death. Studying the linear ubiquitin (linUb) chain assembly complex (LUBAC) was instrumental in proving that immune-receptor-dependent cell death can cause inflammatory disease. As LUBAC regulates both, gene activation and cell death, it can, however, also serve to evaluate the inflammatory-disease-initiating role of gene activation. Investigating mice with two different gene-activation-enhancing point mutations in LUBAC components showed that aberrantly increased gene activation in *Hoil-1^C458A/C458A^*mice did not cause pathology. In contrast, in *Hoip^N101A/N101A^*mice it led to untoward cell death which, unexpectedly, was required for pathological inflammation, culminating in lethal lung fibrosis resembling human idiopathic pulmonary fibrosis (IPF). Hence, we here uncover a new etiology of inflammation, whereby aberrantly enhanced gene activation can be root cause of pathological, fibrogenic inflammation, yet intriguingly via inducing untoward cell death.

## INTRODUCTION

Aberrantly enhanced gene activation was long thought to be the sole driver of tumor necrosis factor (TNF)-induced pathological inflammation^1^. However, this paradigm of inflammatory disease etiology, which had emerged from several genetic mouse models expressing constitutively active NF-κB components^2–6^, changed with the discovery that untoward cell death can be an alternative driver of pathological inflammation^7–10^. This realization predominantly came from the study of the linear ubiquitin (linUb) chain assembly complex (LUBAC), a crucial regulator of the signaling output of many immune receptors^8,11–18^. Whereas LUBAC-mediated linUb is required for efficient receptor-induced gene activation, genetic disruption of LUBAC unexpectedly revealed that its essential physiological function is to suppress pathological cell death. Deletion of either catalytic LUBAC components, HOIL-1 or HOIP, leads to embryonic lethality in mice which depended on RIPK1-kinase-activity and Caspase-8 (CASP8) but is only partially mediated by TNF/TNFR1^7,19^. Similarly, deletion of the adaptor protein SHARPIN, which causes attenuated linUb chain formation, results in a systemic inflammatory syndrome driven by TNFR1-mediated, RIPK1-kinase-activity- and CASP8-dependent apoptosis^8,14,15,20,21^. Intriguingly, humans with inactivating mutations in LUBAC components, despite surviving embryonic development, develop severe systemic autoinflammation and amylopectinosis, of which the auto-inflammatory components can be significantly attenuated with TNF-inhibiting drugs^22–24^. Together, these genetic, functional and pharmacological studies provide compelling evidence that aberrant cell death can initiate inflammatory disease independently of defective inflammatory gene activation.

Despite this conceptual advance, the original hypothesis that excessive inflammatory gene activation itself can drive pathological inflammation has never been directly examined under physiological conditions. Existing evidence supporting this concept relies almost exclusively on constitutive activation of NF-κB signaling^2–6^, rather than enhanced signaling elicited by physiological immune receptor stimulation. Whether selectively increasing stimulation-dependent inflammatory gene activation is sufficient to initiate inflammatory disease therefore remains unknown. Addressing this question is particularly relevant because many autoimmune and autoinflammatory disorders are characterised by aberrantly enhanced gene activation induced by immune receptor stimulation rather than constitutive pathway activation^25^.

LUBAC provides a unique experimental system to address this question because, unlike the core components of the NF-κB signaling machinery, even when constitutively active, it is recruited to receptor signaling complexes only following physiological immune receptor stimulation. Consequently, manipulating LUBAC activity is not expected to activate inflammatory signaling autonomously. Instead, it selectively modulates the magnitude of signaling once immune receptors have been engaged, thereby providing an opportunity to investigate the consequences of enhanced stimulation-dependent, rather than constitutive, inflammatory gene activation *in vivo*^11–18^. To this end, we generated two transgenic models with point mutations in HOIP and HOIL-1, respectively. The underlying scientific rationale was that, under physiological conditions, HOIP-generated linUb chains are negatively regulated by HOIL-1 on the one hand^26^ and by the deubiquitinases (DUBs) CYLD and OTULIN on the other^27–32^. HOIL-1 directly restrains the linUb-generating activity of HOIP by monoubiquitinating LUBAC components^26^, as a point mutation incapacitating the catalytic activity of murine HOIL-1 (C458A) resulted in increased TNF-induced linUb and consequent gene activation^19,26^. Regarding CYLD and OTULIN, it was shown that both DUBs regulate linUb chains, intriguingly however, with opposing functional outcomes^27^. CYLD binds to HOIP via the adaptor protein SPATA2, enabling its recruitment to and subsequent deubiquitination of components of the TNFR1 signaling complex (TNFR1-SC)^33–36^. Thereby, CYLD modulates NF-κB and MAPK signaling downstream of TNFR1. Surprisingly, even though OTULIN forms part of the cytosolic LUBAC via the same interaction site as SPATA2, only CYLD-but not OTULIN-containing LUBAC was found to be recruited to the TNFR1-SC^33,34^. OTULIN inhibits stimulus-responsive LUBAC activity by removing aberrant linear autoubiquitination of its components, thereby preventing stimulation-dependent LUBAC recruitment and consequent LUBAC-dependent gene activation^27,29–32,37^. In line with this, deficiency in OTULIN activity in mice resulted in LUBAC incapacitation and, consequently, aberrant cell death activation^30^. Conversely, mutating the site in human HOIP responsible for LUBAC interaction with both, CYLD-SPATA2 and OTULIN, by a single point mutation, HOIP-N102A, resulted in significantly enhanced linUb in the TNFR1-SC and enhanced TNF-induced gene activation^27^.

Consequently, we here created two transgenic mouse strains, one with a point mutation in the endogenous *Hoil-1* gene (*Hoil-1^C458A/C458A^*, referred to as *C458A*) and another one with a point mutation in the murine *Hoip* gene that corresponded to human HOIP-N102A (*Hoip^N101A/N101A^*, referred to as *N101A*), with the aim to study the potentially pathophysiological consequences of aberrantly enhanced stimulation-dependent gene activation. Unexpectedly, our findings challenge the long-standing view that excessive inflammatory gene activation is, by itself, sufficient to drive inflammatory disease. Instead, we demonstrate that (i) enhanced stimulation-dependent inflammatory gene activation alone is insufficient to initiate pathology; (ii) it becomes pathogenic only when it triggers and thereby converges with aberrant cell death; (iii) this pathological convergence drives spontaneous fibrogenic lung inflammation culminating in lethal pulmonary fibrosis; and (iv) the resulting disease recapitulates multiple histopathological, cellular and molecular features of human idiopathic pulmonary fibrosis (IPF).

## RESULTS

### Enhanced linUb caused by HOIL-1 catalytic inactivation is insufficient to drive spontaneous inflammatory pathology

*C458A* mice were born at the expected Mendelian ratios (Figure 1A, B) and exhibited no overt phenotype throughout their lifespan (up to 18 months of age; Figure 1C). In contrast to previously reported HOIL-1 mutant mice (*Hoil-1^ΔRING1/ΔRING1^*) that developed autoimmunity reminiscent of diseases such as systemic lupus erythematosus (SLE) and Sjögren’s syndrome^38^, *C458A* mice showed no such pathologies. This was consistent with the previously reported *Hoil-1^C458S/C458S^* mice that showed no overt phenotype^39^. Analysis of the *C458A* mice at 18 months of age presented only an increase in different organ weights (Figure 1D) and in serum levels of lactate dehydrogenase (LDH) (Figure 1D). Yet, immunohistochemical (IHC) analysis revealed preserved tissue architecture and only mildly increased immune infiltration in these organs (Figure 1E). Immortalized mouse embryonic fibroblasts (MEFs) derived from *C458A* embryos exhibited a mildly increased susceptibility to TNF-induced cell death (Figure 1F), while TNF stimulation induced a clear increase in linUb chains within the TNFR1-SC (Figure 1G). This surprisingly suggested that enhanced linUb resulting from catalytic inactivation of HOIL-1 does not result in pathological inflammation. Hence, despite aberrantly increased linUb chains in *C458A* mice, the remaining regulatory capacity afforded by CYLD and OTULIN whose interaction with HOIP remains unperturbed in these mice, is sufficient to prevent the development of any overt pathology.

**Figure 1:**
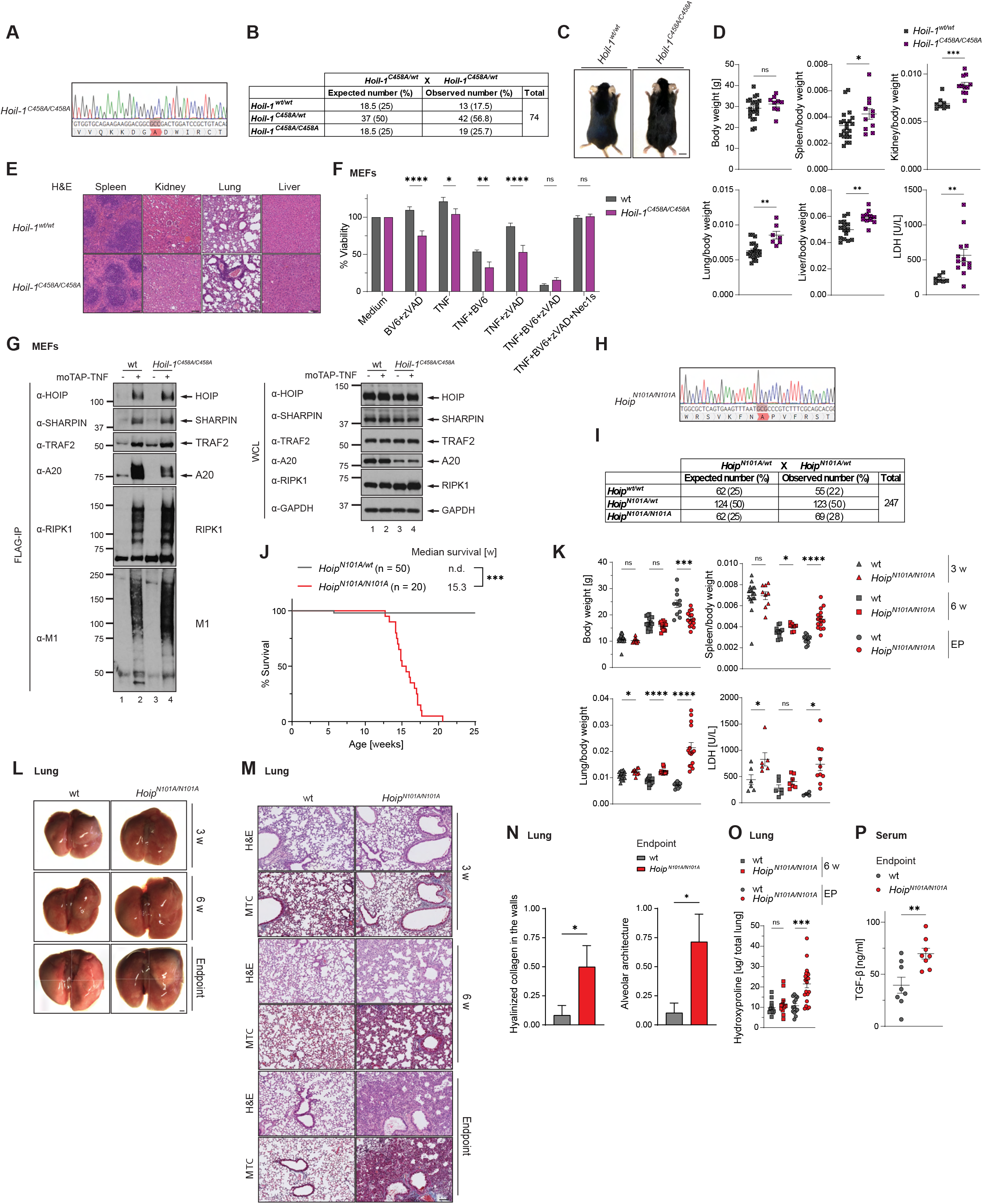
Only *N101A* mice spontaneously develop lethal pulmonary fibrosis despite increased linear ubiquitination in both *N101A* and *C458A* mouse models. (A) Sanger DNA sequencing confirms successful generation of *C458A* mice. (B) Mendelian frequencies obtained from intercrossing *Hoil-1^C458A/wt^* mice. (C) Representative images of male mice with indicated genotype at 18 months of age. Scale bar: 1 cm. (D) Body weight, organ-to-body weight ratio and serum LDH levels for 18-months-old mice of indicated genotypes. Data present mean ± SEM. Statistical analyses were performed via two-tailed unpaired t-test with \**p* < 0.05, \*\**p* < 0.01, \*\*\**p* < 0.001. (E) H&E staining of indicated organs at 18 months of age (n≥7 mice/genotype). Scale bar: 100 μm. (F) MEFs were treated for 24 h as indicated. Cell viability was measured via CellTiter-Glo assay. Data present mean ± SEM (n=9 independent experiments). Statistical analyses were performed via one-way ANOVA with \**p* < 0.05, \*\**p* < 0.01, \*\*\**p* < 0.001, \*\*\*\**p* < 0.0001. (G) MEFs were treated with 1 μg/ml moTAP-TNF for 5 min or left untreated. Lysates were immunoprecipitated with anti-FLAG sepharose beads and analyzed via western blot (n=3). (H) Sanger DNA sequencing confirms successful generation of *N101A* mice. (I) Mendelian frequencies from intercrosses of *Hoip^N101A/wt^* mice. (J) Kaplan-Meier survival analysis of *Hoip^N101A/N101A^* and *Hoip^N101A/wt^* mice. Statistical analysis was performed via Log-rank (Mantel-Cox) test with \*\*\**p* < 0.001. (K) Body weight, organ-to-body weight ratio and serum LDH levels. (L) Representative images of lung. Scale bar: 1 cm. (M) Representative H&E and Masson’s trichrome (MTC) staining of lung (n≥6 mice/group). Scale bar: 100 μm. (N) Pathological evaluation confirms pulmonary fibrosis based on collagen deposition in bronchial walls (0=absent, 1=present) and alveolar architecture integrity (0=normal, 1=distorted, 2=destructed) (n≥7 mice/genotype). (O) Hydroxyproline content in lung homogenates of the indicated genotypes. (P) TGF-β ELISA on mouse serum. Data present mean ± SEM. Statistical analysis was performed via two-tailed unpaired t-test with \**p* < 0.05, \*\**p* < 0.01, \*\*\**p* < 0.001, \*\*\*\**p* < 0.0001. See also Figures S1 and S2.

### Disruption of the LUBAC-DUB interface causes spontaneous lethal pulmonary fibrosis

We and others previously reported that mutation of asparagine 102 (N102) of human HOIP either to alanine (N102A)^27^ or to aspartic acid (N102D)^28^ disrupted its interaction with both, OTULIN and the CYLD-recruiting adaptor protein SPATA2. To compare their respective efficacy in diminishing or possibly abolishing the HOIP–DUB interactions, we performed a biochemical analysis of *Hoip*-deficient A549 cells reconstituted with wild-type (wt) HOIP, HOIP-N102A or HOIP-N102D, so that we could take an informed decision as to which corresponding mutation we should introduce in the mouse model we aimed to develop. This analysis indicated that the interactions with SPATA2 and OTULIN were similarly disrupted in cells expressing HOIP-N102A or HOIP-N102D (Figure S1A). Because we previously showed that in HOIP-N102A-expressing cells TNF-induced linUb and NF-κB activation were increased, we decided to generate the corresponding murine knock-in mutation at position N101 of HOIP, obtaining *Hoip^N101A/N101A^*mice (Figure 1H). This allowed us to directly compare the consequences of enhancing linUb through HOIL-1 catalytic inactivation, where CYLD and OTULIN binding to HOIP remains intact, with those of disrupting the LUBAC-DUB regulatory interface itself. Of note, AlphaFold3-based structural predictions of the LUBAC-DUB complexes followed by FoldX 5.1 energy calculations indicated that both the N102A and N102D mutations of HOIP severely compromised interactions with the two DUBs (Figure S1B–H), yielding predicted ΔΔG values ranging from +4.4 to +6.4 kcal/mol. The differences between the two variants fell within the estimated error margin of the method (∼1 kcal/mol). These findings were further supported by analogous analyses of the corresponding mouse proteins, which produced predicted ΔΔG values ranging from +3.8 to +4.9 kcal/mol (Figure S1I–O).

Analysis of the endogenous LUBAC complex in MEFs derived from *N101A* and wild-type (wt) embryos confirmed that the HOIP-N101A mutation effectively disrupted binding of endogenous OTULIN and CYLD to LUBAC without affecting assembly of the HOIP-HOIL-1-SHARPIN complex (Figure S2A). Consistent with our previous data on human HOIP-N102A, we detected increased linUb at theTNFR1-SC in MEFs obtained from *N101A* as compared to wt embryos as well as increased NF-κB activation, specifically upon TNF stimulation (Figure S2B, C). *N101A* mice were born at normal Mendelian ratios with no embryonic defects (Figure 1I). Intriguingly, however, and in contrast to *C458A* mice, *N101A* mice began to develop overt signs of disease at approximately six weeks of age and succumbed at around 15 weeks, presenting with severe dyspnea and fatigue (Figure 1J). At the survival endpoint, *N101A* mice showed reduced body weight compared to age-matched controls, despite preserved germinal center architecture (Figures 1K and S2E, F), elevated LDH levels and significantly enlarged lungs displaying macroscopic signs of severe organ damage (Figure 1K, L). Histological analysis of multiple organs revealed that the pathological phenotype was highly selective. Skin, intestine and skeletal muscle displayed normal tissue architecture and no increase in immune cell infiltration compared with wt controls (Figure S2G-I). Although the liver and kidneys exhibited reduced organ-to-body-weight ratios, neither organ showed evidence of inflammatory infiltration (Figure S2J-M). In contrast, the heart displayed a significant increase in organ weight accompanied by enhanced immune cell infiltration (Figure S2N, O), most likely secondary to the severe pulmonary disease. Histological examination of the lungs revealed extensive pulmonary fibrosis in *N101A* mice (Figure 1M). Pathological scoring demonstrated marked hyalinised collagen deposition together with profound disruption of alveolar architecture (Figure 1N), while hydroxyproline quantification confirmed a significant increase in collagen content compared with wt lungs (Figure 1O). These findings indicated that *N101A* mice develop histopathological features characteristic of human idiopathic pulmonary fibrosis (IPF)^40^ which we, therefore, investigated further.

Consistent with previous reports in experimental models of pulmonary fibrosis^41^, TGF-β levels were significantly increased in the serum of *N101A* mice (Figure 1P). Finally, hematological analysis provided additional evidence of impaired pulmonary function. *N101A* mice exhibited reduced mean corpuscular volume (MCV) and red cell distribution width (RDW), consistent with chronically impaired blood oxygenation associated with severe lung disease (Figure S2P). In parallel, mean platelet volume (MPV) was significantly reduced, suggesting the presence of smaller, less active platelets and altered platelet homeostasis during chronic pulmonary disease (Figure S2P). Thus, unlike HOIL-1 catalytic inactivation, disruption of the HOIP-DUB interaction results in a selective, lethal and lung-dominant fibrotic disease.

### *N101A* lung disease shows early inflammatory activation, cell death and structural remodeling

To better understand the inflammatory and cellular features of *N101A* fibrotic lung disease, we next performed cytokine profiling of *N101A* lung homogenates. Multiple inflammatory cytokines involved in immune response and extracellular matrix production were significantly elevated in *N101A* mice (Figures 2A and S3A), consistent with enhanced gene-activatory signaling. Several of these cytokines are known to facilitate the recruitment and activation of B cells, monocytes, macrophages, T cells and neutrophils, which secrete pro-fibrotic factors, e.g. TGF-β, and stimulate fibroblast activation as well as extracellular matrix deposition^42^. Strikingly, several of these factors, including CXCL1, IL-33 and CCL5, were already upregulated by 3 weeks of age, implying that pathological processes emerge long before lethality and suggesting a contribution to disease initiation and progression. Notably, some cytokines, including BAFF, CXCL13, S100A9 and CXCL1 were also elevated in the serum of *N101A* mice (Figure S3B) demonstrating that the inflammatory response extended beyond the lung. In line with this, recruitment of immune cells to the lungs of *N101A* mice was already increased at 3 weeks of age (Figure 2B, C). Histological analysis further revealed significantly increased levels of apoptosis and, albeit to a lesser extent, necroptosis within the lungs of *N101A* mice (Figure 2B, C). Together, these observations suggested that the fibrotic disease was unlikely to result solely from enhanced inflammatory gene activation, but rather from the combination of enhanced gene activation with subsequent aberrant cell death.

**Figure 2:**
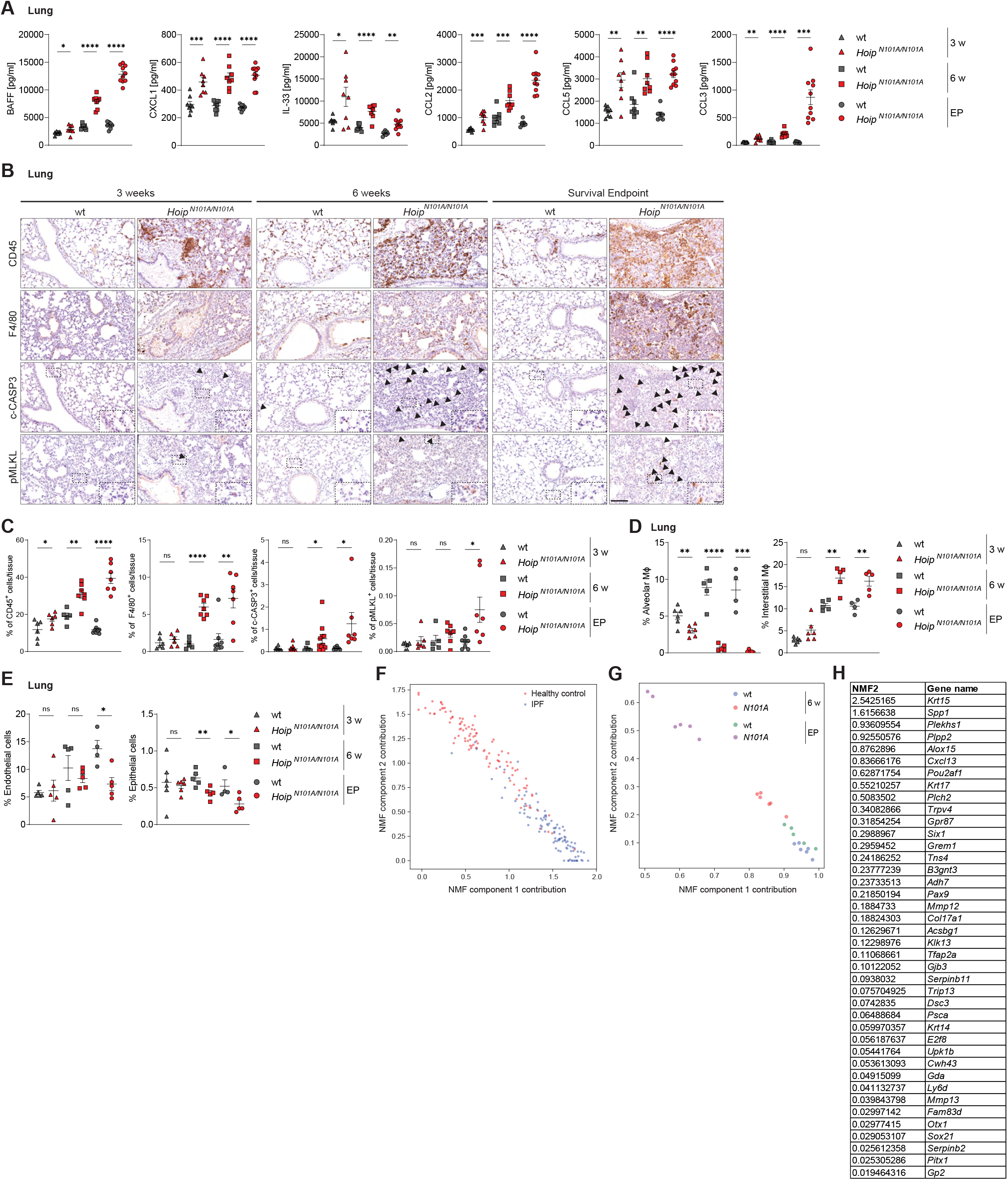
*N101A* mice develop progressive lethal pulmonary fibrosis transcriptionally resembling the human IPF gene expression pattern. (A) Cytokine levels of lung homogenates were analyzed using a Luminex-Multiplex assay for the indicated targets. (B, C) IHC stainings of lungs for CD45, F4/80, cleaved Caspase-3 (c-CASP3) and phospho-MLKL (MLKL pS345) (n≥6 mice/group) with the respective quantifications (C). Scale bar: 100 μm. (D-E) Flow cytometric analysis of lungs at indicated ages. All immune cell subsets represent the percentage from viable CD45^+^ cells. Endothelial and epithelial cells are shown as the percentage of total viable cells. MΦ = macrophages. Data present mean ± SEM. Statistical analyses were performed via two-tailed unpaired t-test with \**p* < 0.05, \*\**p* < 0.01, \*\*\**p* < 0.001, \*\*\*\**p* < 0.0001. (F) Non-negative Matrix Factorization (NMF) analysis of bulk RNA-seq data from human idiopathic pulmonary fibrosis (IPF; n=103) and healthy lung samples (n=103). (G) Translation of human IPF expression patterns to indicated RNA-seq-derived mouse data based on orthology map. (H) List of genes that contribute to IPF according to NMF analysis and are also significantly upregulated in the transcriptome of *N101A* mice at survival endpoint compared to wt controls. The NMF2 values correspond to IPF samples. NMF1 (healthy human lung) is always 0. See also Figures S3, S4, S5 and S10.

Given the central role of immune infiltration and cell death in human pulmonary fibrosis^43^, together with the capacity of infiltrating immune cells to promote extrinsic apoptosis, we next comprehensively characterised the cellular composition of *N101A* lungs and spleens by flow-cytometry across multiple ages.

*N101A* lungs contained reduced levels of CD4^+^ and CD8^+^ T cells as well as NK cells compared with wild-type (wt) mice (Figure S3C). Similarly, CD4^+^ and CD8^+^ T-cell populations were reduced in the spleen (Figure S3D), while histological analysis of the thymus showed increased c-CASP3 staining (Figure S3E, F), suggesting enhanced linUb-dependent T-cell depletion, potentially contributing to systemic inflammation. By contrast, B cells exhibited enhanced survival, consistent with the elevated BAFF levels detected in *N101A* mice (Figure S3B), supporting the development of a B-cell-driven inflammatory and fibrotic phenotype (Figure S3D). Importantly, *N101A* lungs exhibited a marked reduction in alveolar macrophages together with a concomitant increase in interstitial macrophages (Figure 2D), a shift that closely resembles the macrophage composition observed in human IPF patients ^44^. Consistent with the disruption of the endothelial and epithelial barrier that has been linked to epithelial/endothelial-mesenchymal transition (EMT), scarring and fibrosis^45^, both epithelial and endothelial cell populations were significantly reduced in *N101A* lungs (Figure 2E). Collectively, these data demonstrate that HOIP-N101A induces spontaneous lethal pulmonary fibrosis, characterized by pronounced lung inflammation extensive immune remodeling and cell-type-specific cell death.

### *N101A* lung disease recapitulates core transcriptional features of human IPF

Having established that *N101A* mice develop spontaneous pulmonary fibrosis with inflammatory, apoptotic and cellular features relevant to human IPF, we next asked whether this disease also resembles human IPF at the transcriptional level. To do so, we analyzed publicly available bulk RNA-seq data from IPF patients and control samples^46^ and applied Non-negative Matrix Factorization (NMF) analysis to them (see STAR METHODS). The contributions of two key expression patterns effectively separated the IPF and control samples along a spectrum, with each group situated at opposite ends of the two gene expression patterns (Figure 2F). To extend these findings to the *N101A* mouse model, we reduced the expression data to include only genes with clear one-to-one orthologs between human and mouse among the most variable genes (see STAR METHODS). Using this gene orthologous map, we translated the expression patterns derived from human samples to the mouse samples. The human-derived patterns accurately captured the variance in mouse data, clustering *N101A* and wt mice at opposing ends of the NMF contribution spectrum (Figure 2G). Notably, six-week-old *N101A* mice exhibited an intermediate pattern distribution between controls and survival endpoint samples, suggesting that this model mimics a progression of disease at the molecular level. Analysis of the human genes associated with IPF disease revealed 137 transcripts that exclusively contributed to the IPF expression pattern, and not healthy controls. Decisively, 40 of these genes were also significantly upregulated in *N101A* lungs at the survival endpoint (Figure 2H). These results indicate that the *N101A* mouse model mirrors the gene expression variations observed in human IPF patients compared to healthy individuals, confirming the relevance of this model for studying the molecular mechanisms underlying pulmonary fibrosis in patients with IPF. These findings demonstrate that *N101A* lung disease not only shares histopathological and cellular features with human IPF, but also mirrors key elements of the human IPF transcriptional landscape, supporting its relevance for mechanistic dissection of fibrotic lung disease.

### *N101A* lungs show coordinated transcriptomic and proteomic activation of inflammatory, cell death and tissue-remodeling programs

Having established the histopathological, cellular and transcriptional resemblance of *N101A* lung disease to human IPF, we next used multi-omics profiling to define the molecular programs associated with disease initiation and progression. RNA-seq analysis revealed a substantial upregulation of gene transcription in *N101A* lungs as early as at six-weeks of age which was further enhanced at the survival endpoint (Figure S4A, B). Functional annotation using the Hallmark and Gene Ontology Biological Process (GO BP) databases classified these upregulated genes into three main categories: fibrosis, inflammation and cell death (Figure S4C-F). These findings indicated that enhanced linear ubiquitination drives an early transcriptional program converging on these three pathological processes which was already prominently induced at six weeks of age. Specifically, there was early upregulation of key genes involved in the interferon (IFN) response, including transcription factors (e.g., *Irf5*, *Irf7*, *Stat1*, *Stat4*) together with genes involved in nucleic-acid sensing (e.g., *cGas*, *Sting/Tmem173*, *Trex1*, *Ddx60*) (Figure S4G). This inflammatory signature was accompanied by increased expression of pro-inflammatory cytokines such as *Tnf* and *Il1*b as well as chemokines *Cxcl2*, *Ccl5*, *Ccl11*, *Cxcl17* and *Tnfrsf13b* (*BAFF receptor*) (Figure S4H), consistent with the establishment of a robust inflammatory environment capable of recruiting multiple immune cell populations^9^. In parallel, genes involved in programmed cell death were differentially regulated over the course of disease progression. Apoptosis-related genes were already significantly upregulated at six weeks of age, whereas genes associated with necroptosis and pyroptosis displayed a delayed induction, with executioner genes including Mlkl, Gsdme, Casp1 and Ninj1 becoming significantly elevated only at the survival endpoint (Figure S4I). These findings indicate that apoptosis represents the earliest transcriptionally detectable cell death program, while activation of additional lytic cell death pathways accompanies disease progression.

The fibrotic transcriptional program was similarly initiated early during disease development. At six weeks of age, key fibrosis-associated genes involved in extracellular matrix remodeling were already significantly upregulated (Figure S4J). Increased expression of the collagen genes Col1a1, Col5a1, Col5a3 and Col15a1 indicated enhanced extracellular matrix deposition^47,48^, whereas upregulation of *lysyl oxidase* (Lox) suggested increased stabilization of the fibrotic matrix through collagen cross-linking^49^. Elevated expression of Fn1 (fibronectin 1) and Tgfbi (TGF-β-induced protein) indicated enhanced fibroblast activation and adhesion processes^50^, while increased Tgfbr2 expression suggested amplification of the TGF-β signaling pathway, a central driver of fibrosis^51^. By the survival endpoint, these transcriptional changes were further accompanied by robust induction of Wnt10a, indicating activation of Wnt signaling pathways and cooperation with TGF-β signaling to promote fibrogenesis^52^. Additionally, upregulation of keratin genes (*Krt14*, *Krt15*, *Krt17*) and *Gpr87* reflected alterations in the epithelial cell phenotype, potentially indicative of EMT, a process contributing to fibroblast accumulation in fibrotic tissue^53^, whereas elevated S100a4 supported the expansion of activated fibroblasts^54^. Finally, elevated levels of *Alox12e* (*arachidonate 12-lipoxygenase, epidermal-type*) and P4ha3 (*prolyl 4-hydroxylase subunit alpha 3*) suggested enhanced lipid signaling and collagen modification, respectively, both known to facilitate fibrogenesis^55,56^.

Proteomic analyses independently corroborated these transcriptional findings, revealing increased abundance of specific protein clusters also known to be involved in the inflammatory and IFN response, cell death and, specifically, fibrosis (Figure S5A-F). These include STAT3, STAT1, DDX58, EIF6, TMEM173 (STING) and STAT5a/b which were also upregulated at the mRNA level (Figures S4G and S5C). In addition, multiple inflammation-associated proteins were also significantly increased (Figure S5D), reflecting the establishment of a robust inflammatory environment in the lungs of *N101A* mice that may further promote chronic immune-cell recruitment and fibrotic progression^57^. Increased CASP-3 abundance (Figure S5E) corroborated the involvement of apoptosis in disease progression^58^, which was consistent with our histological analyses, confirming a contribution of apoptotic cell death to disease progression.

Proteomic analyses also revealed increased abundance of fibrosis-associated proteins such as EEF2, LDHA, COL14A1, COL6A3, P4HA1, Vimentin (VIM), S100A4, TGFBI, Tenascin C (TNC), and SERPIND1 (Figure S5F), confirming activation of fibrogenic pathways at the protein level. Increased abundance of collagen proteins (COL14A1 and COL6A3), together with enzymes involved in collagen synthesis and modification, including P4HA1 and COLGALT1, indicated enhanced extracellular matrix deposition and remodeling^47,48^. Similarly, elevated levels of TGFBI (TGF-β-induced protein) and S100A4 reflected activated fibroblast phenotypes and sustained TGF-β pathway activation, contributing to fibrosis progression^50,54^, consistent with the transcriptional changes observed (Figure S4J). Thus, transcriptomic and proteomic analyses consistently demonstrate that disruption of the interaction between LUBAC and its regulatory deubiquitinases, CYLD and OTULIN, by the HOIP-N101A mutation induces coordinated activation of inflammatory and cell death pathway components. Together, these data indicate that HOIP-N101A establishes inflammatory and cell death programs that are accompanied by pro-fibrotic tissue-remodeling responses, collectively promoting progressive pulmonary fibrosis.

### Enhanced ubiquitination of fibrotic and inflammatory regulators suggest reinforcement of pro-fibrotic signaling outputs

Given that the HOIP-N101A mutation enhances linear ubiquitination, we next sought to determine how the global ubiquitinome was altered in the lungs of *N101A* mice. DiGLY analysis identified increased ubiquitination of proteins involved in TGF-β signaling and fibrotic processes, including molecular chaperones (HSP90ab1), cytoskeletal components (vimentin, keratin 19, ezrin), metabolic enzymes such as fatty acid synthase (FASN), lactate dehydrogenase A (LDHA), eukaryotic elongation factor 2 (EEF2), and signaling molecules (ICAM1, S100A11, 14-3-3 proteins: YWHAB and YWHAZ) (Figure S6G-I). The coordinated increase in ubiquitination across these proteins suggests that enhanced linear ubiquitination may reinforce multiple pro-fibrotic signaling pathways simultaneously. Specifically, increased ubiquitination of these substrates could stabilize TGF-β receptors and SMAD proteins, thereby amplifying TGF-β signaling, while also influencing EMT through alterations in cytoskeletal dynamics and cell motility, two processes that are central to fibrogenesis. In addition, increased ubiquitination of FASN suggests altered lipid metabolism that may support fibroblast activation and extracellular matrix production, whereas ubiquitination of ICAM1 and S100A11 may enhance immune cell adhesion and fibroblast proliferation, thereby contributing to the inflammatory environment and tissue remodeling characteristic of pulmonary fibrosis.

Ubiquitinome analysis identified enhanced ubiquitination of additional proteins related to inflammation and immune responses, including SAMHD1, AIF1, IIGP1, H2-D1, TMEM176b, PTPRC (CD45), LRBA, IFIT1, NAMPT, GBP2, OAS3, and CD300LD3 (Figure S6J). These findings indicate that dysregulated ubiquitination extends beyond fibrosis-related proteins to encompass key regulators of immune signaling pathways with an established role in modulating and promoting a fibrogenic environment.

Together, these findings demonstrate that disruption of the interaction between LUBAC and its regulatory deubiquitinases, CYLD and OTULIN, amplifies inflammatory signaling at multiple molecular levels. Enhanced linear ubiquitination is accompanied by transcriptional upregulation of genes whose activities have previously been shown to be central to inflammation, cell death and fibrogenesis^47,49,50,52–56^. This relationship is consistently reflected across the transcriptome, proteome and ubiquitinome, with the same signaling pathways and molecular regulators emerging from all three datasets. Beyond promoting inflammatory gene activation, the increased ubiquitination of key signaling proteins further suggests that aberrantly enhanced linear ubiquitination may reinforce and sustain these signaling outputs through the direct or indirect stabilization of non-degradative ubiquitin chains^59^. Together with the coordinated activation of inflammatory signaling detected across the transcriptome, proteome and ubiquitinome, the histological evidence of aberrant cell death and progressive tissue remodeling, and the flow cytometric demonstration of extensive immune-cell recruitment and changes in tissue composition, the data so far support a model in which enhanced inflammatory gene activation and aberrant cell death cooperate to establish a pathological tissue environment that promotes progressive and ultimately irreversible fibrogenesis.

### Disrupted LUBAC-DUB interactions, rather than enhanced linUb alone, drive pulmonary fibrosis

Having established that *N101A* mice develop a spontaneous lung disease displaying transcriptomic, proteomic, ubiquitinomic and histopathological features of pulmonary fibrosis, together with a strong molecular resemblance to human IPF, we next sought to understand why this phenotype was unique to the *N101A* model. Since both the *C458A* and *N101A* mutations enhance linear ubiquitination downstream of immune receptor stimulation, yet only *N101A* mice develop lethal pulmonary fibrosis, we asked whether these distinct pathological outcomes reflect qualitative and/or quantitative differences in LUBAC-mediated signaling activation.

We first compared TNF-induced signaling at the level of the TNFR1-SC. Analysis of this complex in MEFs showed increased linear ubiquitination in both *N101A* and *C458A* cells compared with wt cells (Figure 3D). Although overall linear ubiquitination was modestly higher in *N101A* cells, RIPK1 ubiquitination was more strongly enhanced in *C458A* MEFs (Figure 3D). Despite these differences at the level of individual ubiquitinated substrates, both mutants displayed similarly enhanced TNF-induced gene activation relative to wt cells, as assessed by immunoblotting and RT-qPCR (Figure 3E, F). Thus, the ability of the *N101A* mutation to induce lethal pulmonary fibrosis could not be explained simply by stronger TNF-induced signaling in MEFs.

**Figure 3:**
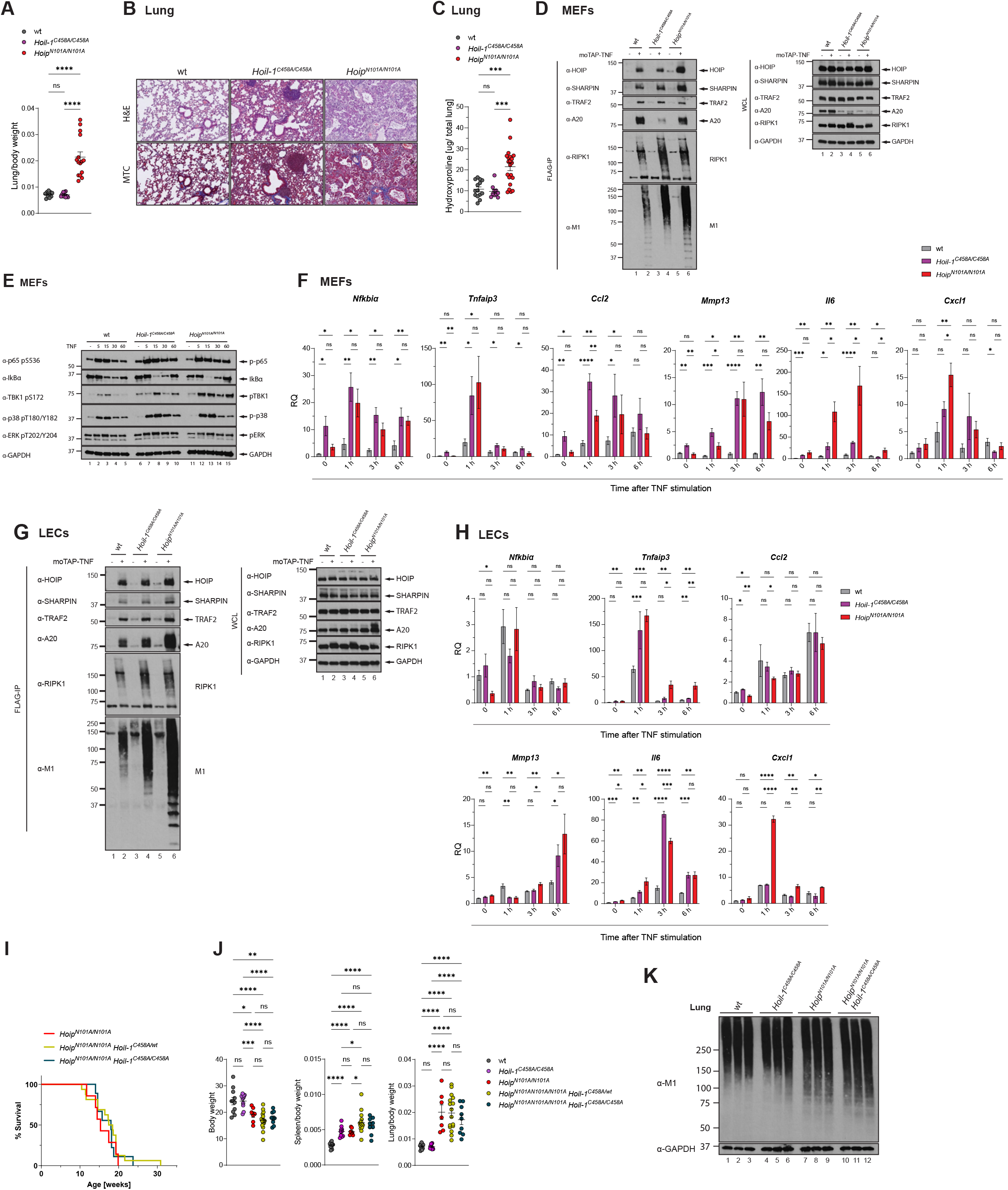
Pulmonary fibrosis results from disrupted LUBAC–DUB interaction rather than from enhanced linear ubiquitination alone. (A, B) Lung-to-body weight ratio (A) and H&E and MTC staining of lung sections (B). Scale bar: 100 μm. All wt and *C458A* mice were sacrificed at around 15 weeks of age, *N101A* and *N101A C458A* mice at their respective humane endpoint. Data present mean ± SEM. Statistical analyses were performed via two-tailed unpaired t-test with \**p* < 0.05, \*\**p* < 0.01, \*\*\**p* < 0.001, \*\*\*\**p* < 0.0001. (C) Hydroxyproline content in lung homogenates of the indicated genotypes. (D) MEFs were treated with 1 μg/ml moTAP-TNF for 5 min or left untreated. Lysates were immunoprecipitated with anti-FLAG sepharose beads and analyzed via western blot (n=3). (E, F) Western blot (E, n=4) and qPCR analysis (F, n=5) of MEFs stimulated with 100 ng/ml TNF for the indicated time points. (G) LECs were treated with 1 μg/ml moTAP-TNF for 5 min or left untreated. Lysates were immunoprecipitated with anti-FLAG sepharose beads and analyzed via western blot (n=3). (H) qPCR analysis of MEFs stimulated with 100 ng/ml TNF for the indicated time points (n=4). qPCR graphs represent relative quantification (log2RQ) normalized to 18S rRNA with expression levels shown relative to untreated wt controls. (I) Kaplan-Meier survival analysis of *N101A C458A* mice and littermate controls. Statistical analysis was performed via Log-rank (Mantel-Cox) test; no significant difference was observed (*p* > 0.05). (J) Body weight and organ-to-body weight ratio. Data present mean ± SEM. Statistical analyses were performed via two-tailed unpaired t-test with \**p* < 0.05, \*\**p* < 0.01, \*\*\**p* < 0.001, \*\*\*\**p* < 0.0001. (K) Western blot of lung homogenates (n=3 biological replicates/genotype). wt and *C458A* mice were sacrificed at around 15 weeks of age, *N101A* genotypes at their respective humane endpoint. See also Figure S6.

Because lung fibrosis in IPF patients is characterized by pulmonary vessel remodeling and hypoxic vasoconstriction leading to vascular dysfunction^40^, we next assessed whether the two mutations differentially affected TNF-induced signaling in lung endothelial cells (LECs). Comparable to the results obtained in MEFs, both *C458A* and *N101A* LECs showed enhanced TNF-induced signaling relative to wt cells, without a qualitative difference that could explain the selective development of fibrosis in *N101A* mice (Figure 3G, H). Together, these data indicate that HOIL-1-C458A and HOIP-N101A similarly potentiate stimulation-dependent TNF-induced gene activation, and that this enhanced gene activation alone is therefore insufficient to account for their divergent in-vivo phenotypes.

We next asked whether the total extent of enhanced linear ubiquitination was responsible for the pulmonary pathology observed in *N101A* mice. To test whether further increasing linear ubiquitination would exacerbate disease, we generated *Hoip^N101/N101A^ Hoil-1^C458A/C458A^*(*N101A C458A*) mice. Decisively, *N101A C458A* animals succumbed with the same kinetics as *N101A* littermate controls and developed comparable pulmonary fibrosis (Figure 3I-K). This suggested that lung fibrogenesis is not determined solely by the amount of enhanced linear ubiquitination. Rather, these findings indicate that disruption of the LUBAC-DUB interaction in *N101A* mice creates a pathological signaling state that is not further aggravated by an additional linUb-enhancing mutation. This identifies the LUBAC-DUB interaction as a critical regulatory node that preserves tissue homeostasis and prevents fibrogenesis *in vivo*.

Having excluded a simple quantitative difference in TNF-induced signaling or total linear ubiquitination as the explanation for the *N101A* phenotype, we next compared the in-vivo consequences of the two mutations in greater detail. Comparative analyses of four-month-old mice revealed that only *N101A* animals developed progressively increasing lung weights accompanied by collagen deposition (Figures 3A-C and S6A, B). By contrast, *C458A* mice did not develop fibrotic lung pathology, but instead showed formation of multiple tertiary lymphoid structures (TLS) throughout the lung parenchyma (Figures 3A-C and S6A, B). Consistent with the absence of overt pulmonary dysfunction, MCV values in *C458A* mice remained comparable to wt controls (Figure S6C). However, peripheral platelet counts were elevated in *C458A* blood, indicative of systemic inflammation (Figure S6C). This interpretation was further supported by increased weights of multiple organs in *C458A* mice, in contrast to *N101A* animals (Figure S6D). *C458A* mice also displayed splenomegaly together with reduced CD4+ and CD8+ T-cell frequencies, resembling the splenic phenotype of *N101A* mice, although in a less pronounced form (Figure S6E-G). In line with systemic inflammation in *C458A* mice, lung homogenates showed similarly increased levels of CCL2, M-CSF and IL-1β in both mutant strains (Figure S6H). However, several cytokines were selectively elevated in *N101A* lungs, including the alarmin IL-33, which is typically released upon epithelial injury (Figure S6H). Consistently, IHC and flow cytometric analyses demonstrated markedly increased immune infiltration in *N101A* lungs (Figure S6I, J). In line with elevated G-CSF levels, granulocytes were selectively expanded in *N101A* lung tissue (Figure S6H, J), a cell population implicated in tissue injury and fibrotic remodeling through the release of reactive oxygen species (ROS) and pro-fibrotic mediators^60–62^. Strikingly, monocyte-derived macrophages, a hallmark immune population in human IPF lungs^63^, were exclusively enriched in *N101A* lungs. These findings indicate that, unlike *C458A*-driven inflammation, the *N101A* inflammatory response is coupled to epithelial injury-associated signals, enhanced immune-cell infiltration, granulocyte expansion and accumulation of monocyte-derived macrophages, thereby establishing a tissue-damaging inflammatory milieu with pro-fibrotic potential. Thus, enhanced linear ubiquitination resulting from HOIL-1 catalytic inactivation was sufficient to induce systemic inflammatory activation and pulmonary lymphoid organization, but not destructive fibrotic lung disease.

The distinct inflammatory and fibrotic microenvironment of *N101A* lungs was further supported by RNA-seq analysis of age-matched wt, *C458A* and *N101A* lungs (Figure S6K-Q). Although several pro-inflammatory and pro-survival pathways were significantly upregulated in both *C458A* and *N101A* lungs, their induction was modestly higher in *N101A* mice. More importantly, however, multiple cell death-associated genes were selectively upregulated in *N101A* lungs, suggesting that the *N101A* mutation does not merely amplify inflammatory gene activation but also establishes a specific transcriptional priming towards cell death.

Together, these findings indicate that enhanced linear ubiquitination caused by removal of the intrinsic negative regulatory brake of LUBAC in *C458A* mice is sufficient to induce systemic inflammatory activation, splenomegaly, altered T-cell frequencies, pulmonary TLS formation and inflammatory cytokine production, but not fibrotic lung pathology. In contrast, disruption of the LUBAC-DUB interface in *N101A* mice does not merely increase stimulation-dependent gene activation beyond that observed in *C458A* mice. Instead, it establishes a qualitatively distinct inflammatory state characterized by epithelial injury-associated cytokines, granulocyte expansion, accumulation of monocyte-derived macrophages, enhanced immune infiltration and selective activation of cell death-associated pathways.

### *N101A* uniquely couples inflammatory gene activation to aberrant cell death in the lung

Having identified selective cell death priming as a defining feature of *N101A* lungs, we next asked which death-inducing pathways and cell types might account for this difference. Because *C458A* mice exhibit enhanced inflammatory signaling and systemic inflammation without developing pulmonary fibrosis, *C458A*-derived cells provided an essential comparator to distinguish cell death responses linked to inflammation alone from those associated with fibrogenic pathology. This allowed us to determine whether *N101A* cells acquire a selective sensitivity to death-inducing stimuli that is absent from a model of enhanced inflammatory gene activation alone. Since the TNF/TNFR1 system has previously been identified as a major pathological driver of inflammation^7,19^, we first assessed TNF-induced cell death in MEFs and LECs derived from wt, *C458A* and *N101A* mice. Because FasL has also been implicated in human IPF^64^, we additionally examined the sensitivity of both mutant models to FasL-induced cell death. *C458A* MEFs displayed a mild increase in sensitivity to TNF-induced cell death, while responding similarly to wt cells following FasL stimulation (Figure 4A, B). By contrast, MEFs derived from *N101A* mice were largely resistant to both TNF- and FasL-induced apoptosis and necroptosis, indicating that disruption of LUBAC-DUB binding prevents death-ligand-induced cell death in fibroblasts (Figure 4A, B). This response was cell-type specific. LECs from *C458A* mice were highly resistant to death ligand-induced cell death (Figure 4C, D). Strikingly, whereas *N101A* LECs responded similarly to wt cells following TNF stimulation, they were selectively vulnerable to FasL-induced apoptosis and necroptosis (Figure 4C, D). Thus, HOIP-N101A does not confer a uniform increase in death-ligand sensitivity across cell types. Instead, it creates a cell-type-specific imbalance in which fibroblasts are protected from death-ligand-induced apoptosis and necroptosis, whereas lung endothelial cells become selectively vulnerable to FasL.

**Figure 4:**
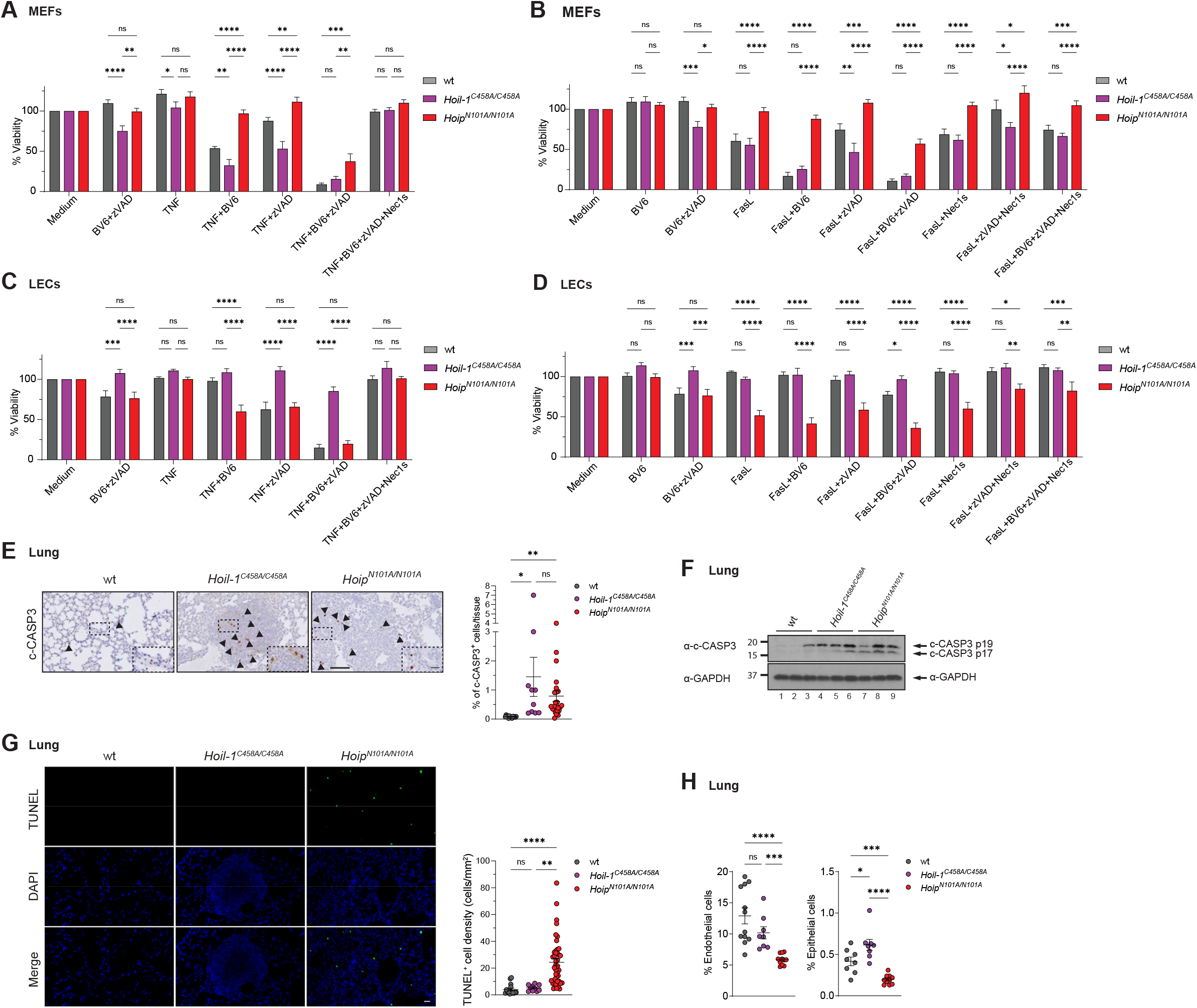
Cell death responses are exclusively increased in *N101A* and not *C458A* lungs. (A-D) MEFs or LECs were treated for 24 h using the indicated agents. Cell viability was measured via CellTiter-Glo assay. Data present mean ± SEM (n≥6 independent experiments respectively). Data present mean ± SEM with *p* values from one-way ANOVA with \**p* < 0.05, \*\**p* < 0.01, \*\*\**p* < 0.001, \*\*\*\**p* < 0.0001. (E) IHC staining of lungs for cleaved Caspase-3 (c-CASP3) with corresponding quantifications. Scale bars: 100 μm and 20 μm. (F) Western blot of lung homogenates (n=3 biological replicates/genotype). wt and *C458A* mice were sacrificed at around 15 weeks of age, *N101A* mice at their respective humane endpoint (median survival = 15 weeks). (G) TUNEL IF staining of lungs with corresponding quantifications. Scale bar: 20 μm. (H) Flow cytometric analysis of lungs from mice of indicated genotypes at 10 weeks of age. Data present mean percentages of total viable cells ± SEM (n ≥ 8 mice per genotype). Statistical analyses were performed via two-tailed unpaired t-test with \**p* < 0.05, \*\**p* < 0.01, \*\*\**p* < 0.001, \*\*\*\**p* < 0.0001. See also Figures S10.

We next examined whether these differential cell death responses were reflected *in vivo*. Although c-CASP3 staining by IHC was similarly increased in both *C458A* and *N101A* lungs (Figure 4E), immunoblot analysis revealed qualitative differences in caspase processing. Specifically, the p19 fragment, which is associated with partial caspase activation, was elevated in *C458A* lungs, whereas the fully active p17 fragment, which mediates irreversible apoptotic execution, was predominantly increased in *N101A* lungs (Figure 4F). Consistently, TUNEL staining demonstrated a significant increase in apoptotic cells exclusively in *N101A* lungs (Figure 4G), indicating that enhanced linUb in the context of preserved HOIP-DUB binding limits full apoptotic execution. In agreement with this interpretation, loss of endothelial and epithelial cells was observed only in *N101A* lungs (Figure 4H). This pattern is consistent with a pro-fibrotic tissue imbalance in which structural cell loss occurs in parallel with relative fibroblast resistance.

Flow-cytometric analyses further supported a FasL-dependent mechanism of structural cell loss in *N101A* lungs. Fas expression was significantly increased on *N101A* epithelial cells compared with wt and *C458A* mice, whereas the mild increase in Fas expression on *C458A* epithelial cells likely reflected the inflammatory milieu present in these animals (Figure S6R). Moreover, FasL expression was significantly elevated across multiple immune cell populations in *N101A* lungs, potentially favoring FasL-dependent killing in this genetic background (Figure S6R).

Thus, although *C458A* mice develop systemic inflammation, their preserved control of death-ligand-induced cell death may allow tissue adaptation without progression to destructive fibrotic pathology. By contrast, *N101A* mice combine enhanced linUb-driven inflammatory gene activation with aberrant, cell-type-specific death regulation: fibroblasts are protected from death-ligand-induced apoptosis and necroptosis, whereas epithelial and endothelial cells become selectively vulnerable, particularly to FasL. This imbalance provides a mechanistic explanation for how enhanced inflammatory signaling becomes coupled to structural cell loss, tissue remodeling and progressive fibrotic lung disease in *N101A* mice.

### TNFR1 and Fas contribute to lung inflammation and cell death in *N101A* mice

Although the ex-vivo analyses revealed differential sensitivity of *N101A* cells to TNF- and FasL-induced cell death, isolated cell-type-specific assays cannot fully recapitulate the multicellular complexity of pulmonary fibrosis. *In vivo*, death receptor signaling occurs within an inflammatory tissue environment in which immune cells, epithelial cells, endothelial cells and stromal populations interact and collectively shape tissue injury, repair and remodeling. Therefore, to determine the contribution of TNFR1 and Fas signaling to the development of autochthonous fibrotic lung disease in *N101A* mice, we genetically deleted either *Tnfr1* or *Fas* in the *N101A* background by crossing *N101A* mice with *Tnfr1*^-/-^ and *Fas*^-/-^ mice, respectively.

Co-deletion of TNFR1 delayed fibrosis onset and provided a modest, but significant, survival advantage to *N101A* mice (Figure 5A). However, at the survival endpoint, body weight and lung weight were not significantly different between *N101A* and *N101A Tnfr1^-/-^* mice (Figures 5B and S7A-C). By contrast, spleen weight and serum LDH levels were significantly reduced in *Tnfr1*-deficient *N101A* mice compared with *Tnfr1*-proficient *N101A* controls, suggesting reduced systemic inflammation and tissue damage (Figure 5B). Consistent with the unchanged lung weights, Masson’s trichrome staining and hydroxyproline quantification confirmed comparable fibrosis development between the genotypes (Figures 5C and S7D).

**Figure 5:**
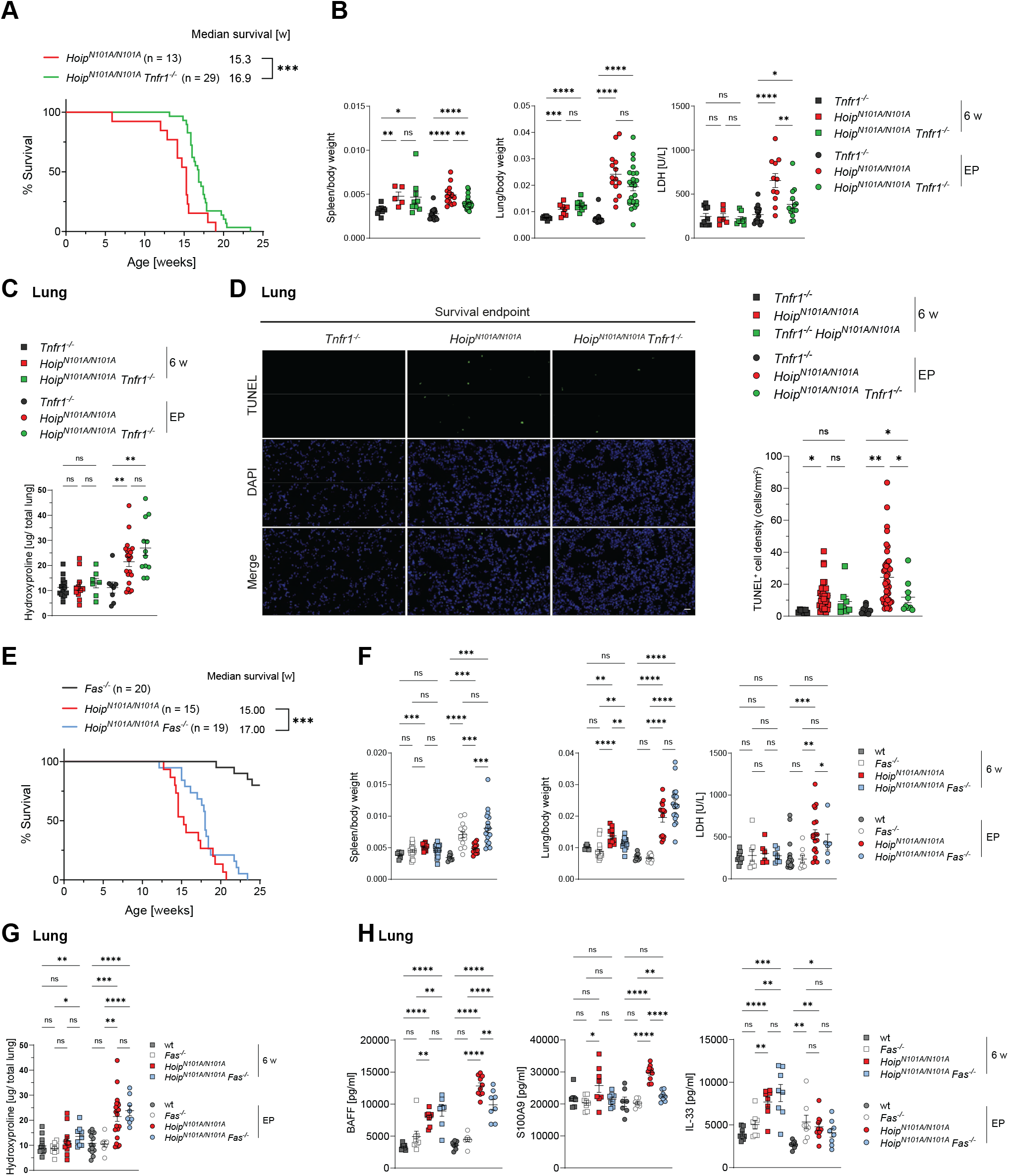
TNFR1 and Fas contribute to lung inflammation and cell death in *N101A* mice. (A) Kaplan-Meier survival analysis of *Hoip^N101A/N101A^ Tnfr1^-/-^* compared to *Hoip^N101A/N101A^* mice. Statistical analysis was performed via Log-rank (Mantel-Cox) test with \*\*\**p* < 0.001. (B) Organ-to-body weight ratio and serum LDH levels. (C) Hydroxyproline content in lung homogenates of the indicated genotypes. (D) TUNEL IF staining of lungs with corresponding quantifications. Scale bar: 20 μm. (E) Kaplan-Meier survival analysis of *Hoip^N101A/N101A^ Fas^-/-^* compared to *Hoip^N101A/N101A^* mice. Statistical analysis was performed via Log-rank (Mantel-Cox) test with \*\*\**p* < 0.001. (F) Organ-to-body weight ratio and serum LDH levels. (G) Hydroxyproline content in lung homogenates of the indicated genotypes. (H) Cytokine levels of lung homogenates were analyzed using a Luminex-Multiplex assay for the indicated targets. Data present mean ± SEM. Statistical analysis was performed via two-tailed unpaired t-test with \**p* < 0.05, \*\**p* < 0.01, \*\*\**p* < 0.001, \*\*\*\**p* < 0.0001. See also Figures S7 and S8.

Despite the similar extent of fibrosis at endpoint, TNFR1 deficiency altered specific pathological features of the *N101A* phenotype. Inflammatory cytokine levels and immune infiltration were largely comparable between *N101A* and *N101A Tnfr1^-/-^* lungs (Figure S7E, F), whereas cell death levels were significantly reduced upon TNFR1 deficiency at the survival endpoint (Figures 5D and S7F). These findings indicate that TNFR1-mediated signaling contributes to cell death, systemic inflammation and disease kinetics in *N101A* mice. Because TNFR1 deficiency reduced cell death *in vivo* despite the absence of pronounced TNF sensitivity in the ex vivo fibroblast and LEC assays, TNFR1-dependent cell death may occur in additional lung-resident or infiltrating cell populations. Consistent with previous studies showing that TNF inhibitors provide only mild benefit in patients with IPF^65,66^, these data suggest that TNFR1 contributes to disease severity and tissue damage but is not the dominant etiological driver of *N101A*-mediated fibrogenesis. This distinguishes the *N101A* model from OTULIN-related autoinflammatory syndrome (ORAS), in which TNF inhibition alone can provide substantial benefit^67^, and suggests that simultaneous disruption of HOIP binding to both OTULIN and CYLD creates a broader inflammatory and cell death-driven pathological state.

We next analyzed the contribution of Fas signaling in *N101A Fas^-/-^* mice. Loss of Fas is known to cause a lymphoproliferative disorder^68,69^ associated with premature death at approximately 30 weeks of age in mice. Fas deficiency prolonged survival of *N101A* mice; however, similar to *N101A Tnfr1^-/-^* animals, *N101A Fas^-/-^* mice ultimately succumbed only around two weeks later than *N101A* controls (Figures 5E and S8A). Splenomegaly in *N101A Fas^-/-^* mice was comparable to that observed in age-matched *Fas^-/-^* controls (Figures 5F and S8B). Lung weight and the extent of fibrosis were also unchanged across all *N101A* genotypes (Figures 5F, G and S8C, D). As observed upon Tnfr1 loss, however, Fas deficiency reduced serum LDH levels in *N101A* mice, suggesting diminished tissue damage (Figure 5F).

Luminex multiplex profiling further revealed that more than half of the measured inflammatory cytokines and chemokines were significantly reduced in *N101A* lungs upon Fas loss (Figures 5H and S8E). Surprisingly, the alarmin IL-33 was also elevated in four-month-old *Fas^-/-^* mice relative to wt controls, suggesting ongoing epithelial stress in *Fas*-deficient mice independently of HOIP-N101A expression. Consistent with this, immune infiltration was increased in *Fas*-deficient lungs (Figure S8F), indicating that Fas deficiency alone can drive a basal inflammatory state. As F4/80 levels were unaffected, this suggests that, owing to the underlying lymphoproliferative disorder, immune cell populations distinct from those accumulating in *N101A* lungs infiltrate the lung in *Fas*-deficient mice.

Together, these findings demonstrate that both TNFR1 and Fas contribute to distinct aspects of the *N101A* lung phenotype, but neither pathway alone is sufficient to account for fibrogenesis. TNFR1 deficiency reduces cell death, systemic inflammation and disease kinetics without preventing fibrosis, whereas Fas deficiency improves survival and ameliorates the inflammatory cytokine profile but its contribution is confounded by the intrinsic lymphoproliferative and lung-inflammatory phenotype of *Fas*-deficient mice. Thus, death receptor signaling contributes to lung inflammation and tissue damage in *N101A* mice, but the persistence of fibrosis upon deletion of either *Tnfr1* or *Fas* indicates that *N101A*-driven fibrogenesis is not dependent on a single death receptor pathway.

### Combined attenuation of apoptosis and necroptosis delays *N101A* disease progression

Considering the increased apoptosis detected in *N101A* lungs, together with the partial amelioration of disease-associated features upon genetic deletion of either *Tnfr1* or *Fas*, we next sought to determine whether CASP8 contributes to *N101A*-driven lung pathology. CASP8 represents the apical initiator caspase commonly engaged downstream of multiple death receptor pathways, including TNFR1 and Fas, and therefore provided a convergent genetic entry point to assess the contribution of death receptor-induced apoptosis to fibrotic disease in *N101A* mice. However, complete deletion of Casp8 is embryonically lethal^70^. Although co-deletion of *Casp8* together with *Mlkl* or *Ripk3* prevents this lethality, homozygous *Casp8/Mlkl*- or *Casp8/Ripk3*-deficient mice are not ideal to dissect the respective contributions of CASP8-dependent apoptosis and RIPK3/MLKL-dependent necroptosis to *N101A*-driven fibrosis. This is because these mice develop lymphoproliferative syndromes, similar to *Fas^-/-^* mice, at around 8 and 14 weeks of age, respectively^70,71^, thus overlapping with the time window of fibrotic disease development in *N101A* mice. Moreover, because *Ripk3* is located in close chromosomal proximity to *Hoip* on mouse chromosome 14, it is not possible to generate *N101A Ripk3^-/-^* mice by conventional genetic crossing.

Given that heterozygous loss of *Casp8* is sufficient to alleviate TNFR1-driven dermatitis in *cpdm* mice^8^, we hypothesized that CASP8 haploinsufficiency might similarly ameliorate lung disease in *N101A* mice. We therefore assessed the contribution of apoptosis and necroptosis by combining Casp8 heterozygosity with genetic deletion of *Mlkl*. Whereas loss of *Mlkl* or *Casp8* heterozygosity alone had no effect on disease progression, combined targeting of both pathways significantly prolonged the survival of *N101A* mice (Figure 6A). Nevertheless, all animals ultimately developed pulmonary fibrosis and succumbed to dyspnea, with lung pathology indistinguishable from that of *N101A* controls (Figure 6B-F). Immune-cell infiltration was likewise comparable across genotypes at the humane endpoint (Figure S9A). Consistent with this, unchanged levels of c-CASP3 staining and TUNEL positivity indicated that Casp8 haploinsufficiency was insufficient to suppress apoptotic execution at late stages of disease (Figures 6G and S9A).

**Figure 6:**
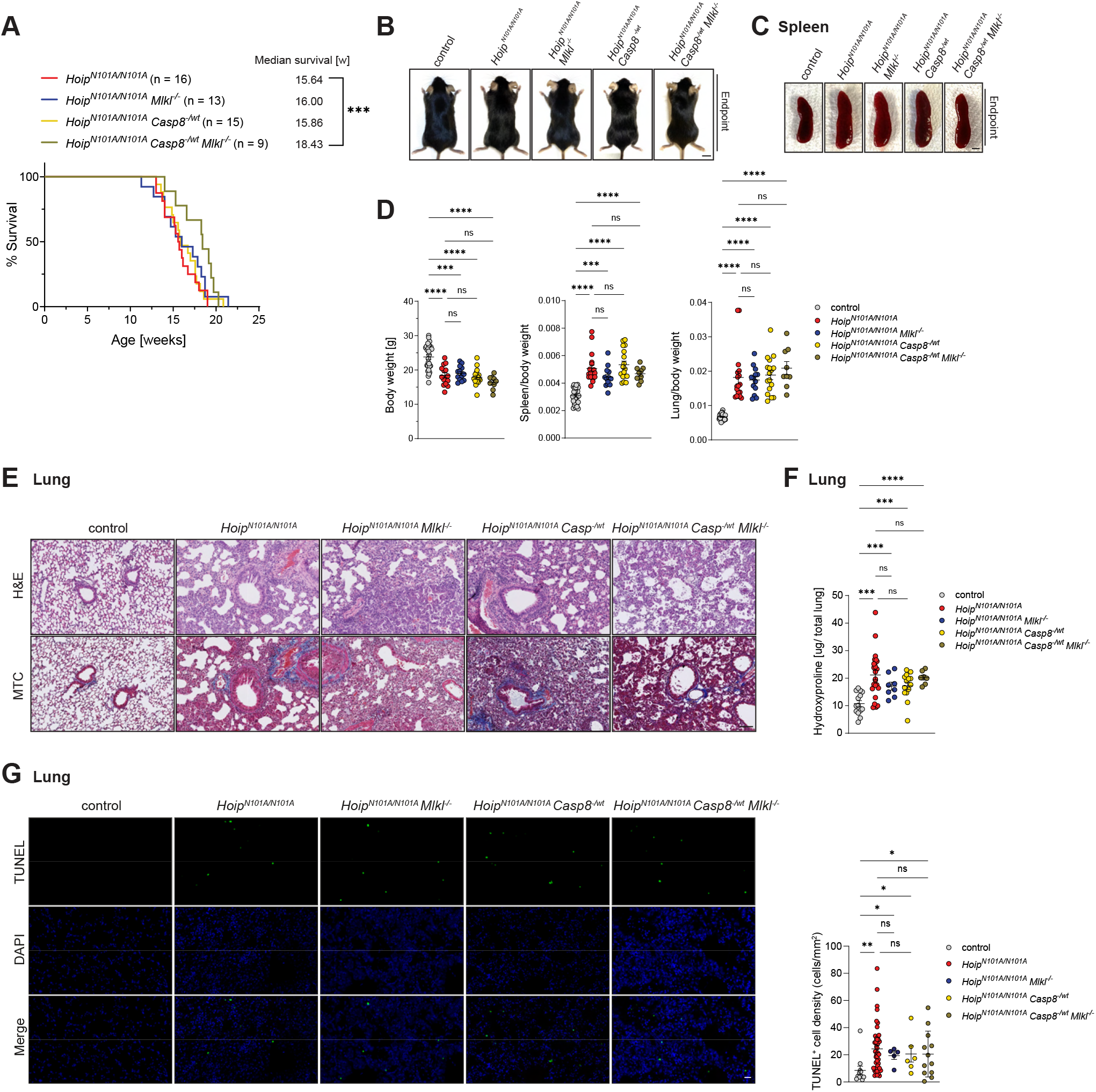
Concomitant *Mlkl* deficiency and heterozygous *Casp8* loss prolongs the survival of *N101A* mice. (A) Kaplan-Meier survival analysis of indicated genotypes. Statistical analysis was performed via Log-rank (Mantel-Cox) test with \*\*\**p* < 0.001. (B, C) Representative images of male mice (B) and spleens (C) with indicated genotypes. Scale bar: 1 cm and 200 μm respectively. (D) Body weight and organ-to-body weight ratio. (E) Representative H&E and MTC staining of lung (n≥6 mice/group). Scale bar: 100 μm. (F) Hydroxyproline content in lung homogenates of the indicated genotypes. (G) TUNEL IF staining of lungs with corresponding quantifications. Scale bar: 20 μm. Data present mean ± SEM. Statistical analysis was performed via two-tailed unpaired t-test with \**p* < 0.05, \*\**p* < 0.01, \*\*\**p* < 0.001, \*\*\*\**p* < 0.0001. See also Figure S9.

These results support a model in which CASP8-dependent apoptosis is functionally involved in *N101A*-driven disease progression, with necroptosis contributing secondarily to disease progression, while incomplete targeting of apoptotic execution is unable to halt established fibrogenesis. However, because Casp8 was only reduced by haploinsufficiency, these experiments do not fully eliminate apoptotic execution and therefore cannot exclude a stronger requirement for apoptosis under conditions of complete pathway blockade. Together, these findings indicate that apoptotic and necroptotic pathways contribute to disease kinetics, but that partial reduction of apoptosis, even in the absence of MLKL-dependent necroptosis, is insufficient to prevent established fibrotic lung pathology.

### Combined TNF and FasL blockade delays lethal disease progression in *N101A* mice

The genetic analyses showed that deletion of either *Tnfr1* or *Fas* alone partially ameliorated selected features of the *N101A* phenotype but was insufficient to prevent lethal pulmonary fibrosis (Figure 5). We therefore hypothesized that simultaneous inhibition of TNF- and FasL-dependent signaling might provide greater therapeutic benefit by targeting two death receptor pathways that contribute to lung inflammation, tissue damage and disease progression. To avoid the intrinsic lymphoproliferative phenotype associated with germline Fas deficiency, we treated *N101A* mice from four weeks of age with TNF or FasL inhibitors, either individually or in combination, using TNFR2-Fc and Fas-Fc (Figures 7A and S9B). Control mice received mIgG2a antibodies to account for potential Fc-mediated effects. The use of Fas-Fc was further supported by recent studies in mice infected with MA20, a mouse-adapted SARS-CoV-2 strain, in which FasL blockade reduced lung cell death and inflammation and extended survival^72^. In *N101A* mice, neither TNFR2-Fc nor Fas-Fc alone significantly affected survival. By contrast, combined TNFR2-Fc and Fas-Fc therapy significantly prolonged survival (Figures 7B and S9C), indicating that simultaneous inhibition of both pathways provides therapeutic benefit in this model. Although the reduction in lung weight at the humane endpoint did not reach statistical significance, spleen weight was significantly decreased in *N101A* mice treated with TNFR2-Fc alone or with combined TNFR2-Fc and Fas-Fc therapy (Figures 7C and S9D), consistent with a systemic anti-inflammatory effect and in line with the genetic models.

**Figure 7:**
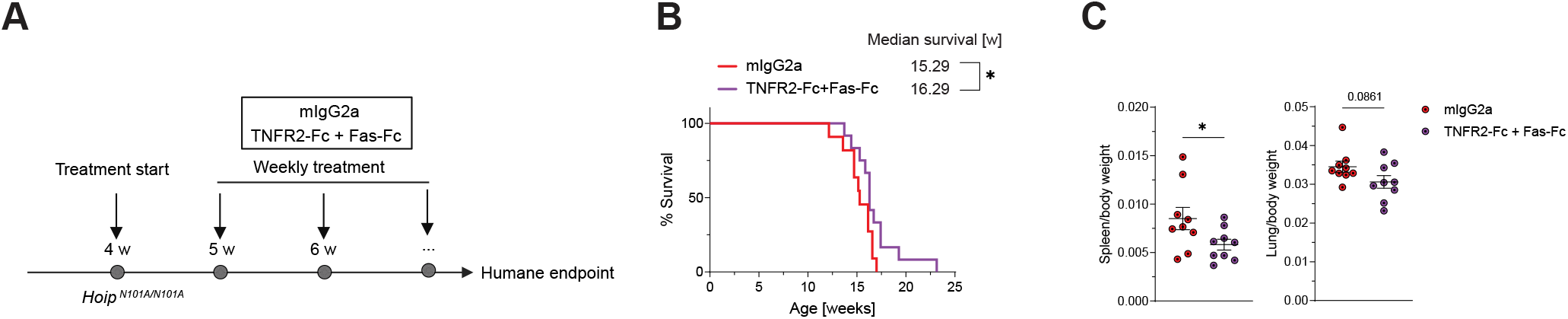
Combined inhibition of TNF and FasL provides therapeutic benefit in HOIP-N101A-driven lung fibrosis. (A) Experimental design: 4-week-old *N101A* mice were randomized into the indicated treatment groups. 500 µg of each compound was administered intraperitoneally once per week from 4 weeks of age until the humane endpoint. (B) Kaplan-Meier survival analysis of *N101A* mice treated with the indicated Fc proteins. Statistical analysis was performed via Log-rank (Mantel-Cox) test with \**p* < 0.05. (C) Organ-to-body-weight ratios. Data present mean ± SEM. Statistical analysis was performed via two-tailed unpaired t-test with \**p* < 0.05. See also Figure S9.

Together, these findings indicate that TNF and FasL signaling make partially overlapping, non-redundant contributions to the inflammatory and tissue-damaging phenotype of *N101A* mice. While blockade of either pathway alone is insufficient to alter disease outcome, combined pharmacological inhibition delays lethal disease progression. These data provide proof-of-principle that therapeutic targeting of convergent death receptor pathways can ameliorate fibrotic lung disease driven by the pathological coupling of inflammatory gene activation and aberrant cell death, and suggest that modulation of cell death-associated inflammatory pathways warrants further investigation in human IPF.

## DISCUSSION

In this study, we identify HOIP-N101A mice as a genetically defined autochthonous model of spontaneous, progressive and lethal pulmonary fibrosis. This discovery emerged from an effort to interrogate the pathological consequences of enhanced LUBAC-dependent linear ubiquitination under conditions of physiological immune receptor stimulation. By generating two knock-in mouse models in which linear ubiquitination is enhanced through distinct mechanisms, we found that HOIL-1 catalytic inactivation, which increases stimulation-dependent linear ubiquitination while preserving HOIP binding to CYLD and OTULIN, does not cause overt inflammatory pathology. By contrast, disruption of the HOIP interaction with both CYLD and OTULIN leads to severe fibrotic lung disease. Thus, increased linear ubiquitination and enhanced stimulation-dependent inflammatory gene activation are not, by themselves, sufficient to drive pathological inflammation. Rather, our data show that inflammatory gene activation becomes pathogenic when it converges with aberrant cell death, thereby creating a tissue-damaging inflammatory environment that promotes progressive and ultimately irreversible fibrogenesis. These findings have important implications for how the etiology of pathological inflammation is understood. Aberrantly enhanced gene activation has long been considered a major driver of inflammatory disease, a view strongly influenced by seminal studies identifying NF-κB as a central regulator of inflammatory gene expression^2–6^. This concept was later expanded by the discovery that untoward cell death can itself initiate lethal inflammation^7^. As a result, inflammation-driven diseases are now generally viewed as arising either from aberrant gene activation or from excessive cell death^73–76^. However, whereas the genetic evidence supporting untoward cell death as a driver of inflammatory disease is now substantial, evidence that enhanced gene activation alone can initiate inflammatory pathology has largely relied on models of constitutive NF-κB activation^2–6^. Whether increased stimulation-dependent gene activation, as occurs downstream of physiological immune receptor engagement, is sufficient to cause inflammatory disease has remained unresolved.

LUBAC provided a unique genetic system to address this question because it regulates both gene-activatory and cell-death-controlling outputs downstream of immune receptors. By generating two mouse models with enhanced LUBAC-dependent linear ubiquitination through mechanistically distinct routes, we were able to separate enhanced stimulation-dependent gene activation from disruption of LUBAC-DUB-mediated tissue protection. The *C458A* model showed that increased linear ubiquitination and enhanced stimulation-dependent gene activation caused by HOIL-1 catalytic inactivation are not sufficient to induce spontaneous inflammatory pathology. By contrast, disruption of the HOIP interaction with both OTULIN and CYLD in *N101A* mice caused progressive and lethal pulmonary fibrosis. These findings challenge the idea that enhanced inflammatory gene activation is, by itself, sufficient to drive inflammatory disease. Instead, they support a model in which enhanced gene activation becomes pathogenic when it converges with aberrant cell death, thereby creating a tissue-damaging inflammatory environment that promotes fibrogenesis. Thus, the LUBAC-DUB interface emerges as an essential regulatory node that preserves tissue homeostasis by restraining both excessive inflammatory signaling and pathological cell death *in vivo*.

A major limitation of many currently available preclinical fibrosis models is that they rely on acute chemical injury, most commonly bleomycin, or on forced activation of individual pro-fibrotic pathways^77–82^. Although these models have been invaluable, they often fail to fully recapitulate the chronic, progressive and complex nature of human IPF. *N101A* mice, by contrast, develop spontaneous, progressive and invariably lethal pulmonary fibrosis. This disease is accompanied by early inflammatory cytokine induction, immune-cell remodeling, epithelial and endothelial cell loss, apoptotic cell death, accumulation of interstitial and monocyte-derived macrophages, and activation of pro-fibrotic tissue-remodeling programs. Importantly, *N101A* lung disease also recapitulates core transcriptional features of human IPF, with human IPF-derived gene expression patterns separating *N101A* and control mouse lungs and positioning six-week-old *N101A* mice between healthy controls and endpoint disease. These findings support the *N101A* mouse as an autochthonous model of progressive fibrotic lung disease with molecular and cellular relevance to human IPF. The comparison between *C458A* and *N101A* mice was particularly informative because both models exhibit enhanced LUBAC-dependent signaling, yet only *N101A* mice develop fibrotic lung disease. This demonstrates that the extent of enhanced linear ubiquitination or TNF-induced gene activation alone cannot explain fibrogenesis. Instead, *N101A* mice develop a qualitatively distinct inflammatory state characterized by epithelial injury-associated cytokines, granulocyte expansion, accumulation of monocyte-derived macrophages and selective transcriptional priming of cell death pathways. Ex-vivo and in-vivo analyses further revealed a cell-type-specific imbalance in death responsiveness: fibroblasts were relatively protected from death ligand-induced apoptosis and necroptosis, whereas lung endothelial and epithelial compartments showed evidence of FasL-associated vulnerability and structural cell loss. Such an imbalance provides a plausible mechanistic link between enhanced inflammatory signaling, loss of tissue integrity and progressive fibrotic remodeling. Our genetic analyses further indicate that TNFR1 and Fas contribute to *N101A* disease but are not individually required for fibrogenesis. Deletion of *Tnfr1* delayed disease onset, reduced systemic inflammation and diminished cell death, yet failed to prevent pulmonary fibrosis. This is consistent with the limited clinical benefit of TNF inhibition in patients with IPF^65,66^ and distinguishes the *N101A* model from OTULIN-related autoinflammatory syndrome (ORAS), in which TNF blockade can provide substantial therapeutic benefit^67^. Fas deficiency also prolonged survival and reduced inflammatory cytokine levels in *N101A* lungs, but interpretation of this experiment was complicated by the intrinsic lymphoproliferative and lung-inflammatory phenotype of *Fas*-deficient mice^68,69^. Together, these findings suggest that *N101A*-driven fibrosis is not dependent on a single death receptor pathway, but instead reflects a broader pathological state in which multiple death receptor and inflammatory signals contribute to tissue damage and disease progression. Consistent with this interpretation, partial genetic attenuation of apoptosis and necroptosis by combining *Casp8* haploinsufficiency with *Mlkl* deletion significantly prolonged survival, whereas targeting either pathway alone was insufficient. Nevertheless, combined genetic attenuation did not prevent fibrosis, indicating that partial reduction of CASP8-dependent apoptosis, even in the absence of MLKL-mediated necroptosis, is insufficient to halt disease progression once the *N101A* pathological program is established. These data support a functional contribution of apoptotic and necroptotic pathways to disease kinetics, while also highlighting that complete suppression of the relevant cell death program was not achieved genetically. Thus, the *N101A* model suggests that cell death contributes to fibrotic disease not as a single isolated pathway, but as part of a broader inflammatory tissue circuit that reinforces structural damage and remodeling.

The therapeutic experiments further support this concept. While inhibition of TNF or FasL alone did not significantly alter survival, combined TNFR2-Fc and Fas-Fc treatment delayed lethal disease progression and reduced systemic inflammation. These findings provide proof-of-principle that simultaneous targeting of convergent death receptor pathways can ameliorate disease in a model where fibrogenesis emerges from the pathological coupling of inflammatory gene activation and aberrant cell death. Importantly, these data should not be interpreted as evidence that TNF and FasL are the only relevant death receptor pathways in fibrotic lung disease. Rather, they indicate that death receptor-associated inflammatory and cell death circuits can be therapeutically modulated and warrant further investigation in the context of IPF and related fibrotic disorders.

Collectively, this work identifies a previously unrecognized etiology of inflammatory fibrotic disease in which enhanced stimulation-dependent inflammatory gene activation becomes pathogenic only when coupled to aberrant cell death. In the *N101A* model, disruption of the LUBAC-DUB interface converts enhanced receptor-induced signaling into a tissue-damaging inflammatory program characterized by immune-cell remodeling, structural cell loss and progressive fibrogenesis. These findings provide mechanistic insight into how inflammatory signaling and cell death can cooperate to drive irreversible lung fibrosis and establish the *N101A* mouse as a genetically defined model to dissect the molecular logic of fibrotic disease. More broadly, they raise the possibility that similar convergence between inflammatory gene activation and aberrant cell death may contribute to fibrotic pathology also when caused, e.g., by interferon and/or in other organs, providing a rationale for exploring cell death-associated inflammatory pathways as therapeutic targets in human fibrotic disease.

## ACKNOWLEDGMENTS

We would like to thank the members of the Liccardi and Walczak laboratories as well as Dr. Nieves Peltzer for helpful discussions. In addition, we would like to thank the Histology Core Facility of SFB1403, the CECAD *in vivo* Research Facility staff of the University of Cologne and the Cologne Center for Genomics for their support. This study was funded by the Alexander von Humboldt Foundation (H.W.), a Wellcome Trust Investigator Award (214342/Z/18/Z to H.W.), a Medical Research Council Grant (MR/S00811X/1 to H.W.), a Cancer Research UK Programme Grant (A27323 to H.W.), the Deutsche Forschungsgemeinschaft through SFB1399 Project C06 (413326622) (H.W., G.L.), SFB1530 Project A03 (455784452) (H.W., G.L.) and SFB1403 Project A12 (414786233) (H.W.), the CANcer TARgeting (CANTAR) Network grant by the Ministry of Art and Science of the State of NRW, Germany (H.W., G.L.), the Institute of Biochemistry I, Medical Faculty, University of Cologne (G.L.), “BIOSMALL” EU (HORIZON-MSCA-2022-SE-01 project to B.D.), the Hungarian National Research, Development and Innovation Office (TKP2021-EGA-33, FK-143751 and FK-147045 to B.D. and Z.M.), Bolyai Research Scholarship of the Hungarian Academy of Sciences (Z.M.) and the Association for the Study of Lung Cancer / International Lung Cancer Foundation Young Investigator Grant 2022 (Z.M.).

## AUTHOR CONTRIBUTIONS

Conceptualization: H.W., G.L.

Supervision: H.W., G.L.

Animal licenses and documentation: J.S., E.R.

Experimental design: J.S., H.W., G.L.

Preliminary experimental work: J.S., H.R.

In-vivo experiments: J.S., D.B. (Figure 1B-F), C.T. (Figures 7 and S9B-D)

Generation of *Fas^-/-^*mice and FasL production: S.S.S., J.S.

Biochemical experiments: J.S.

IHC and IF stainings: J.S., C.K., K.K. (Figure 2B)

DiGLY proteomics: L.S., M.K., J.S.

AlphaFold 3 predictions: L.G., P.E., J.S., G.L.

Pathological scoring: Z.M., B.D.

Design and analysis of flow cytometry data: J.S., A.M.

RNA-seq analysis including NMF: N.A., J.S., G.L.

Visualization: J.S., G.L.

Writing: J.S., H.W., G.L.

## RESOURCE AVAILABILITY

### Lead contact

Further information and reasonable requests for reagents may be directed to, and will be fulfilled by, the lead contact Henning Walczak.

### Materials availability

All unique reagents generated in this study are available upon reasonable request from the lead contact.

### Data and code availability

The mass spectrometry proteomics data have been deposited to the ProteomeXchange Consortium via the PRIDE partner repository with the dataset identifier PXD057938. The analyzed RNA-seq data of human IPF patients are publicly available via the Gene Expression Omnibus database: GSE150910. All other data are available in the main text or the supplementary materials. Any additional information required to reanalyze the data reported in this paper is available from the lead contact upon reasonable request.

## STAR METHODS

### Mice

*C458A* mice were generated under mixed C57BL/6N and C57BL6/J background with Cyagen Biosciences. All other mouse strains were bred on a clean C57BL/6N background. *N101A* and *Fas^-/-^* mice were generated via CRISPR/Cas9-mediated gene editing in the Transgenic Core Unit (TCU) in the *in vivo* Research Facility (ivRF) of the CECAD Research Center of the University of Cologne. Knockout of *Fas* was achieved via insertion of a premature stop codon in exon 2. *Tnfr1^-/-^*, *Mlkl^-/-^* and *Casp8^-/wt^* mice were obtained from the SFB1403 mouse repository at the CECAD Research Center. All in-vivo experiments were conducted with mixed genders. The mice were housed in individually ventilated cages under a stable 12 h day/night cycle at the animal facilities of the CECAD Research Center and provided water and food *ad libitum* under stable room temperature (RT) and humidity. Mice were kept under specific pathogen free (SPF) conditions with pathogen testing on a quarterly basis. Animals were monitored daily and sacrificed via cervical dislocation once endpoint criteria were reached. All experiments were performed according to German animal protection laws and approved by local ethics committees as well as local government authorities (*Landesamt für Natur, Umwelt und Verbraucherschutz Nordrhein Westfalen* under 81-02.04.2020.A022 and 2024.800).

### In-vivo treatments

Murine TNFR2-Fc and Fas-Fc were produced by WuXi Biologics as previously described^72^. Four-week-old mice were randomized to treatment groups, with both sexes represented. Recombinant Fc proteins (500 µg) were administered intraperitoneally once weekly in 200 µL PBS. PBS alone served as the vehicle control. Treatment continued until humane endpoint criteria were reached. All animal procedures were approved and performed under project licence PP6471236.

### Tissue processing and immunohistochemistry

Tissues were fixed in 4% paraformaldehyde (Walter CMP GmbH, WAL60622) for 24 h prior to embedding in paraffin using a vacuum tissue processor (Leica Biosystems, ASP200S) through standard automatic tissue embedding protocols. The paraffin-embedded blocks were sectioned into 3 µm (intestine), 4 µm (lung) or 5 µm (heart, kidney, muscle, liver, skin and spleen) consecutive cuts, transferred on polysine adhesion slides and dried at 40°C overnight. Lungs were stained via Masson’s trichrome (Sigma-Aldrich, HT15) or like the other organs via hematoxylin-eosin (Sigma-Aldrich, HT1079). After de-waxing and rehydration, the antigen retrieval for CD45 (550539, BD Biosciences, 1:200), F4/80 (MCA497R, Bio-Rad, 1:100), and c-CASP3 (9661, Cell Signaling Technology, 1:200) staining were performed in sodium citrate buffer, pH 6 (C9999, Sigma-Aldrich) for 10 min at 114°-121°C using the TintoRetriever Pressure Cooker. Slides for c-CASP3 (9664, Cell Signaling Technology, 1:50) staining in skin were retrieved in sodium citrate buffer, pH 6 in a water bath at 60-65°C overnight. MLKL pS345 staining was performed as reported ^83^. After the retrieval, sections were blocked using Bloxall (SP-6000-100, Vector Laboratories) and Avidin/Biotin blocking kit (SP-2001, Vector Laboratories). Primary antibodies were incubated overnight at 4°C followed by incubation with biotinylated secondary goat anti-rat antibody (EC-BA-9400, Biozol, 1:200) or goat anti-rabbit (VEC-BA-1000, Biozol, 1:200). IHC signal was obtained using the Vectastain Elit ABC-HRP Detection Kit (PK-6100, Vector Laboratories) and DAB substrate (SK-4105, Vector Laboratories). For F4/80 staining, ImmPRESS HRP Goat Anti-Rat (MP-7444-15, Vector Laboratories) secondary antibody and amplification with the TSA Biotin Systems (NEL700A001KT, Akoya Biosystems) was applied before the ABC-HRP kit. After counterstaining with hematoxylin (H-3401, Vector Laboratories), sections were rehydrated and mounted using mounting media (H-5700, Vector Laboratories or P36930, Thermo Fisher Scientific). Tissue sections were digitalized using the Nanozoomer S360 (Hamamatsu) slide scanner. Quantification was performed blinded either with the whole tissue or indicated sections using the positive cell detection toll of QuPath^84^. False-positive signals (e.g. dust, folded tissue areas, shadows) were excluded from the analysis.

### TUNEL immunofluorescence staining

Terminal deoxynucleotidyl transferase dUTP Nick End Labeling (TUNEL) assay was performed according to the manufacturer’s instructions (Promega, G3250), with the exception of the Proteinase K digestion step. Briefly, deparaffinized tissue sections were fixed in 4% formaldehyde in PBS for 15 min at RT, followed by two washes in PBS. Sections were permeabilized with Proteinase K (20 µg/ml) for 15 min at RT, then washed in PBS and post-fixed in 4% formaldehyde for 5 min. After washing, sections were equilibrated in Equilibration Buffer for 10 min. The Terminal deoxynucleotidyl Transferase (TdT) reaction mix was applied to the tissue, covered with plastic coverslips, and incubated for 60 min at 37 °C in a humidified chamber protected from light. The reaction was terminated by immersion in 2x saline-sodium citrate (SSC) for 15 min, followed by three PBS washes. Afterwards, slides were counterstained with DAPI (Sigma Aldrich, D9564) for 10 min and washed three times with PBS. Slides were mounted with Invitrogen™ ProLong™ Gold Antifade Mountant (Thermo Fisher Scientific, P36930). Fluorescent signals were acquired using a SLIDEVIEW VS200 Universal Whole Slide Imaging Scanner (Evident) and analyzed on QuPath.

### Pathological scoring

Lungs were scored for fibrosis features in a blinded fashion. 3 independent foci per lung with n≥7 mice/genotype were scored for two criteria: Hyalinized collagen deposition in the bronchial walls (0=absent, 1=present) and alveolar architecture integrity (0=normal, 1=distorted, 2=destructed).

### AlphaFold3 structure prediction and FoldX energy calculations

LUBAC-DUB complex structures were predicted using AlphaFold3. For each complex, the complete LUBAC ternary complex (HOIP, HOIL-1L, and SHARPIN) was modeled together with either OTULIN or SPATA2 and CYLD, using canonical UniProt sequences for both human (HOIP: Q96EP0; HOIL-1L: Q9BYM8; SHARPIN: Q9H0F6; OTULIN: Q96BN8; SPATA2: Q9UM82; CYLD: Q9NQC7) and mouse (HOIP: Q924T7; HOIL-1L: Q9WUB0; SHARPIN: Q91WA6; OTULIN: Q3UCV8; SPATA2: Q8K004; CYLD: Q80TQ2) orthologs. Predictions were performed on the AlphaFold3 server (alphafoldserver.com) with a model seed of 42. The top-ranked models were converted from CIF to PDB format using UCSF ChimeraX 1.8. To estimate the effect of HOIP-N102 (human) and HOIP-N101 (mouse) mutations on complex stability, FoldX 5.1 BuildModel calculations were performed. Structures were first energy-minimized using the FoldX RepairPDB function to optimize side-chain conformations. The mutations N102A, N102D (human) and N101A, N101D (mouse) were then introduced individually, and the change in Gibbs free energy (ddG, kcal/mol) was calculated using default parameters (298 K, pH 7, ionic strength 0.05 M). Positive ddG values indicate destabilization of the complex upon mutation.

### Cell culture

Primary mouse embryonic fibroblasts (MEFs) were isolated from embryos at E13.5. Minced embryos were incubated in 0.5% Trypsin/EDTA (P10-024100, PAN-Biotech) at 37°C for 10 min. Digested embryos were passed through a 40 μm filter and transferred to a 10cm dish (Corning, 35003) in DMEM (P04-03550, PAN-Biotech) with 4.5 g/L glucose, 20% FBS (Sigma-Aldrich, F7524) and 1% Penicillin/Streptomycin (PAN-Biotech, P06-07100).

Lung endothelial cells (LECs) were isolated from 8- to 12-weeks-old mice. Lungs were minced and incubated with 1 ml collagenase II (Sigma-Aldrich, C2-22) at 37°C for 45 min. Tissue suspensions were smashed through a 70 μm cell strainer and centrifuged at 4°C and 500xg for 5 min. The pellet was resuspended in Buffer 1 (PBS, 0.5% FBS, 2 mM EDTA) and incubated with 30 μl of CD31-coated beads (Miltenyi Biotec, 130-097-418) at 4°C for 15 min. Afterwards, the cell suspension was centrifuged again and resuspended in 500 μl Buffer 1. Samples were run through LS columns (Miltenyi Biotec, 130-042-401) and flushed using a plunger. Directly after primary cell isolation, cells were cultured in endothelial cell medium constituted of DMEM (Gibco, 11960044) with 4.5 g/L glucose, 20% FBS, 2 mM Glutamine (Gibco, 25030081), 1% Penicillin/Streptomycin, 1 mM sodium pyruvate (Gibco, 11360070), HEPES (Gibco, 12440061) and 1% non-essential amino acids (PAN-Biotech, P08-32100) for six days. Afterwards cells were cultured in a 1:1 mix of endothelial cell medium and endothelial cell growth medium (PromoCell, C-22011).

Primary MEFs and LECs were immortalized via SV40 LT infection. For this, pBABE-puro SV40 LT vector was packaged into Platinum-E cells. Viral supernatant and 10 µg/ml polybrene (Sigma-Aldrich, TR-1003-G) were added to each primary cell culture. Spin infection was performed via centrifugation at 2.500 rpm and 32°C for 40 min. Immortalized cells were selected with 1 µg/ml puromycin for 7 days.

*Hoip*-deficient A549 cells have been previously generated^27^ and transiently transfected as described^33^. PEI was used as transfection reagent.

### Cell viability assay

1 x 10^4^ cells were seeded per 96-well and incubated at 37°C overnight. On the next day, cells were treated as indicated: 100 ng/ml recombinant moTAP-TNF (self-produced), 20 µM Z-VAD-FMK (APExBIO, APE-A1902), 2 µM BV6 (Selleck Chemicals, S7597), 10 µM Nec-1s (Selleck Chemicals, S8251), 10 ng/ml recombinant His-izFasL (self-produced). 24 h after treatment, a CellTiter-Glo^®^ Luminescent Cell Viability Assay (Promega, G7573) was performed. Luminescence was measured in a Spark^®^ Multimode Microplate Reader (TECAN).

### Protein lysis and immunoprecipitation

5 x 10^4^ cells or 8 x 10^6^ cells were seeded into 6-wells (Greiner Bio-One, 657160) or 15cm dishes (Thermo Scientific, 168381), respectively and incubated at 37°C overnight. On the next day, cells were treated with 100 ng/ml or 1 µg/ml recombinant moTAP-TNF (self-produced). After the treatment, cells for TNFR complex I immunoprecipitation (IP) were lysed in lysis buffer 1 (30 mM Tris-HCl (pH 7.5), 120 mM NaCl, 2 mM EDTA, 2 mM KCl, 10% glycerol, 1% Triton X-100), while cells for whole-cell-lysates were lysed in lysis buffer 2 (20 mM Tris-HCl (pH 7.5), 150 mM NaCl, 2 mM EDTA, 10% glycerol, 1% Triton X-100). Both lysis buffers were supplemented with PhosSTOP (Roche, 4906845001), cOmplete™ EDTA-free Protease Inhibitor Cocktail (Roche, 11873580001) and 50 µM PR-619 (Selleckchem, S7130). Protein lysates were centrifuged for 15 min at maximum speed (Eppendorf, 5427 R) and transferred into fresh Eppendorf tubes. Protein concentrations were measured via Pierce™ BCA® Protein Assay (Thermo Scientific, 23227). For the input lysate, protein samples were mixed with 1X NuPAGE™ LDS Sample Buffer (Invitrogen, NP0008) and boiled at 95°C for 10 min. Samples for TNFR complex I IP were incubated with 30 µl anti-FLAG® M2 Affinity beads (Sigma-Aldrich, A2220) at 4°C overnight. For SHARPIN-IP, 3 µg/ml anti-SHARPIN antibody (Proteintech, 14626-1-AP) was incubated with 20 µl Protein G Sepharose^TM^ 4 Fast Flow beads (Cytiva, 17061801) at 4°C for 4 h. Afterwards, beads were washed with lysis buffer 2 and added to the protein lysates. IPs were incubated at 4°C overnight and washed five times with lysis buffer 2 on the next day. Samples were boiled in 2X NuPAGE™ LDS Sample Buffer at 95°C for 10 min.

### Lung homogenate preparation

30 mg of lung tissue were homogenized in 300 µl RIPA buffer (Thermo Fisher Scientific, 89901) supplemented with PhosSTOP (Roche, 4906845001), cOmplete™ EDTA-free Protease Inhibitor Cocktail (Roche, 11873580001) and 50 µM PR-619 (Selleckchem, S7130) at 30 Hz for 2 min in 3 cycles using stainless steel beads (Qiagen, 69989) and the TissueLyser II (Qiagen). Subsequently, lung homogenates were incubated for 1 h on ice, centrifuged for 15 min at maximum speed and transferred into fresh Eppendorf tubes. Further processing was performed as described above for cell protein lysates.

### Western blot analysis

Protein lysates were loaded onto Criterion TGX Stain-Free Precast Gels (Bio-Rad, 5678085) and transferred on Trans-Blot Turbo Midi 0.2 µm Nitrocellulose (Bio-Rad, 1704159). Membranes were blocked with 5% Bovine Serum Albumin (BSA; Sigma-Aldrich, A8022) in 1X Tris-Buffered Saline with Tween 20 (TBST) for 1 h and subsequently incubated in primary antibody (Table S2) at RT overnight. All primary antibodies were diluted 1:1,000 in 5% BSA in 1X TBST. After washing the membranes for 30 min in 1X TBST, they were incubated in secondary antibody (α-rabbit IgG: Jackson ImmunoResearch, 211-032-171; α-mouse IgG: Jackson ImmunoResearch, 115-035-174; α-rat IgG: Jackson ImmunoResearch, 112-035-175; α-sheep IgG: Millipore, AP184P) for 1.5 h. All secondary antibodies were diluted 1:5,000 in 5% powdered milk (Carl Roth, T145.3) in 1X TBST. Following the incubation, membranes were washed and signal detection was performed using Enhanced Chemiluminescence (ECL) reagents (PerkinElmer, NEL105001EA). The signal was visualized by exposing the membranes to X-ray films (Valmex, 807947).

### LDH measurement

Serum lactate dehydrogenase (LDH) was measured with a standard assay in a Cobas C 702 biochemical analyzer (Roche).

### Hydroxyproline quantification

Hydroxyproline assay was performed according to manufacturer’s protocol (Sigma, MAK008). In brief, 10 mg of lung tissue were homogenized in 100 µl water at 30 Hz for 2 min in 2 cycles using stainless steel beads (Qiagen, 69989) and the TissueLyser II (Qiagen). Lung homogenates were centrifuged at 10,000 rcf for 1 min. The supernatant was mixed 1:1 with 37% hydrochloric acid in a screw vial (Glastechnik Gräfenroda GmbH, GHS6-10R-SKFW32-H). The samples were hydrolyzed at 120°C for 3 h. Afterwards, samples were centrifuged at 10,000 rcf for 3 min and 10 µl of supernatant transferred to a 96-well plate (Thermo Fisher Scientific, 442404). Samples and hydroxyproline standard were dried at 60°C in an oven. Then, all samples as well as standards were oxidized by adding 100 µL of chloramine T/oxidation buffer mixture and incubating at RT for 5 min. Subsequently, 100 µL of diluted 4-(dimethylamino) benzaldehyde (DMAB) reagent was added and the reaction was incubated for 90 min at 60 °C to allow color development. Absorbance was then measured at 560 nm in a TECAN.

### Cytokine analysis

Lungs were snap-frozen in liquid nitrogen and homogenized in cell lysis buffer 2 (R&D systems, 895347) supplemented with PhosSTOP (Roche, 4906845001), cOmplete™ EDTA-free Protease Inhibitor Cocktail (Roche, 11873580001) and Benzonase® Nuclease (Millipore, 70746-4) using the TissueLyser II (Qiagen). Lung homogenates were incubated for 1 h at −80°C, thawed on ice and subsequently centrifuged for 15 min at maximum speed (Eppendorf, 5427 R). Supernatants were transferred into fresh Eppendorf tubes and protein concentrations measured via Pierce™ BCA® Protein Assay (Thermo Scientific, 23227). Cytokine analysis on lung homogenates and serum was conducted using a customized Luminex® Multiplex assay (R&D Systems) according to manufacturer’s instruction. TGF-β levels were quantified by ELISA (R&D Systems, DY1679) following the supplier’s guidelines.

### Hematological analysis

Blood was collected from the vena cava and promptly transferred to EDTA tubes (Sarstedt, 20.1278). Differential blood count was performed with 50 µl peripheral blood using the VETSCAN® HM5 according to manufacturer’s instructions.

### Flow cytometry

Chopped lungs were incubated in 1.5 ml Liberase/DNase solution buffer (0.3 mg/ml Liberase TL, Roche, 5401119001; 0.2 mg/ml recombinant DNase I, Roche, 4716728001) at 37°C for 30 min. Spleens and digested lungs were passed through a 70 µM strainer (Greiner, 542070) and centrifuged at 1.500 rpm and 4°C for 5 min. Red blood cell lysis was performed with 1X RBC lysis buffer (BioLegend, 420302). The reaction was stopped after 4 min with PBS and the samples centrifuged again. Spleen and lung cells were blocked with CD16/32 (1:100; BioLegend, 101320) for 30 min at 4°C, washed with FACS buffer (PBS + 1% FBS + 5 mM EDTA) and stained with viability dye (1:1,000; Thermo Scientific, 65-0865-18). After 30 min, samples were washed and incubated with the respective antibodies (Table S3) in Brilliant Stain Buffer (BD, 566349) for 30 min at 4°C. After washing, Strep antibody (1:100; BioLegend, 405225) was incubated for 30 min at 4°C. Samples were measured on a BD FACSymphony™ A3 Cell Analyzer. Data were analyzed using FlowJo™ v10.10 and GraphPad Prism 10. The gating strategy is illustrated in figure S9.

### 3’ mRNA-sequencing

RNA was isolated from lung tissue using the RNeasy Plus Kit (Qiagen, 74136) according to the manufacturer’s protocol. For 3’ mRNA sequencing, 2 µg of RNA was submitted to the Cologne Center for Genomics (CCG), where library preparation was carried out using the Lexogen QuantSeq 3’ mRNA sequencing protocol. The quality and quantity of the library were validated and quantified using TapeStation and qPCR, respectively. Sequencing was performed on a NovaSeq 6000 platform using a single-read (SR) 1×100 bp sequencing protocol.

### Data processing and analysis of 3’ mRNA-seq for *N101A* and *C458A* lung comparison

Raw sequencing reads were processed by GeneVia Technologies using the following workflow. Reads were processed using fastp (v0.23) for quality control and adapter trimming. Filtered reads were aligned to the Mus musculus reference genome (GRCm38) using STAR (v2.7), and transcript-level quantification was performed in alignment-based mode using Salmon (v1.10.2). Quality control metrics were aggregated with MultiQC (v1.21). Differential gene expression analysis was conducted using DESeq2 (v1.46.0) with the design formula ∼ Condition. Genes with fewer than five total counts across all samples were excluded. Differentially expressed genes (DEGs) were defined as those with an adjusted p-value <0.05 and absolute log2 fold change >1, with multiple testing correction performed using the Benjamini-Hochberg method. Gene annotations were retrieved via biomaRt (v2.62.1) with Ensembl annotations. Pathway enrichment was performed using gene set enrichment analysis (GSEA) against MSigDB Hallmark and Gene Ontology Biological Process (GOBP) gene sets, implemented in clusterProfiler (v4.14.6). Genes were ranked by log2 fold change, and gene sets containing between 15 and 500 genes were considered. Visualizations were generated using Python (v3.8.12) with Matplotlib (v3.7.5) and Seaborn (v0.13.2).

### Data processing and analysis of 3’ mRNA-seq from 6-week-old and EP *N101A* datasets

Sequencing reads were trimmed by cutadapt tool (version 4.1) and mapped to murine reference transcriptome GRCm38/mm10 using Start aligner (version 2.7.0e). Read counts were estimated using RSEM tool (version 1.3.1). The RSEM gene level Counts were used as expression values. Protein coding genes with sum Counts over samples less than 10 were removed. Count values were normalized to 10^4^ across all genes and then log transformed (ln(TPM+1)) for heatmap plots and PCA analysis. In case of heatmaps, expression values were further transformed to z-scores. Differential gene expression analysis was carried out in python using the pyDESeq2 (version 0.4.3) with default parameters. The integer part of the raw RSEM gen level counts were used as the input of pyDESeq2. The gseapy python package (version 1.1.0) applied to differential gene expression data to determine enrichment of the 50 MSigDB hallmark gene sets (2023.1.Mm). Pre-ranked gene lists were generated by combining the DESeq2 stat value. All parameters were left to their default values.

### Expression Pattern Analysis

Gene expression patterns and their corresponding contributions in each sample were identified from RNA sequencing data using Non-negative Matrix Factorization (NMF), following the previously described method^85^. This approach reduces the dimensionality of transcriptomic profiles by assuming a limited number of expression patterns and simultaneously determining the contribution of each pattern in each sample’s transcription profile. In this process, the data, which originally included thousands of gene expressions, was reduced to a smaller number of expression patterns and their contributions in each sample. These patterns were represented as linear combinations of gene expressions, with specific relative ratios. The NMF calculations were performed using the “decomposition. NMF” module from the “sklearn” (version 1.3.1) Python package, with the default “Coordinate Descent” solver and the ‘random’ initialization parameter on a macOS platform.

We extracted k patterns based on the expression of N genes across a sample cohort of size M. Model selection in NMF, which involved determining the optimal number of patterns k and evaluating the model’s robustness, followed the method as previously described^85^. For each k in in 2..11, Nr=250 NMF runs were performed with different random initializations. During each run, samples were clustered into k groups based on their highest-expressed patterns. The pattern with the highest contribution in a sample determined the cluster assignment of that sample.

A Connectivity Matrix (C) was created, where each element c_i,j was 1 if samples i and j were co-clustered otherwise they would correspond to a 0 if they did not. The Consensus Clustering Matrix was calculated as the average of the Connectivity Matrices across all Nr NMF runs. The Cophenetic Correlation Coefficient ρ^86^ was used to assess clustering robustness across different runs. The coefficient, which ranges from 0 to 1, measures the correlation between the hierarchical linkage of the samples based on the matrix C and the Euclidean distance of matrix 1-C. A higher ρ value indicates more stable clustering. The k with the highest ρ was selected as the optimal number of expression patterns (k=2). The analysis of ρ for various k values was conducted using human RNA-seq data, given the availability of a large dataset. The “scipy” (version 1.13.1) Python package was used to compute the Cophenetic Correlation Coefficient.

### Comparison of gene expression patterns in human and mouse

For both human and mouse, the top 5,000 highly variable genes were selected. Of these, 4,256 human genes and 4,252 mouse genes were identified as one-to-one orthologs based on annotations from the Ensembl database. However, 1,912 genes were common between the two sets, and these were used for our comparative analysis. In the human dataset, gene expression patterns and their contributions for each sample were derived using the NMF approach, based on the expression of the one-to-one homologous gene set. Through homology relationships, the human expression patterns were translated to mouse. The contributions for the mouse data were then calculated by linearly regressing the translated human patterns onto the mouse expression data.

### Proteomic analysis and ubiquitin diGly proteomics

For the diGly-remnant study, lungs from mice sacrificed at six-weeks of age or survival endpoint were collected and directly frozen in liquid nitrogen. Organs were homogenized in cell lysis buffer 2 (R&D Systems, 895347) supplemented with PhosSTOP (Roche, 4906845001), cOmplete™ EDTA-free Protease Inhibitor Cocktail (Roche, 11873580001), Benzonase® Nuclease (Millipore, 70746-4) and 100 µM PR-619 (Selleckchem, S7130) using the TissueLyser II (Qiagen). Debris was removed by centrifugation at maximum speed (Eppendorf, 5427 R) and 4°C for 15 min. Protein concentration was determined using Pierce™ BCA® Protein Assay (Thermo Scientific, 23227). Tissue lysates were reduced and alkylated with final concentrations of 5 mM TCEP and 15 mM 2-chloroacetamid pH 7.5 (Sigma-Aldrich) for 45 min at RT with constant shaking at 750 rpm. Protein digestion was performed following the SP3 protocol ^87^. In brief, both hydrophilic and hydrophobic beads were added to the sample and bound by adding 1:1 volume of acetonitrile (ACN). After 8-minutes incubation time, magnetic beads were immobilized and washed 2x with 70% ethanol and acetonitrile. Proteins were digested with trypsin (substrate:enzyme ratio 100:1) overnight at RT. Samples were acidified with TFA (pH < 2) and desalted using Sep-Pak C18 1 cc Vac Cartidge, 50 mg (Waters, Milford, US). Eluate was dried using a speed vac concentrator. 10 µg were resuspended in 5% formic acid, 2% acetonitrile.

Immunoaffinity Purification (IAP) with diGLY antibody was performed with PTMScan Ubiquitin Remnant Motif (K-ε-GG) (Cell Signaling, 5562) and manufacturer’s instructions were followed. In brief, desalted peptides were reconstituted in 1 ml 1X IAP buffer. Antibody beads were washed four times with 1 ml of 1X PBS and centrifuged at 2,000 x g after each wash. Beads were resuspended in PBS and 40 µl were added to the desalted peptides. Samples were incubated on a rotator for 2 h at 4°C, followed by centrifugation at 2,000 x g for 30 sec. Beads were washed two times with 1 ml 1X IAP and three times with HPLC water. Peptides were eluted two times with 55 µl 0.15% TFA. Samples were cleaned up using Stop-and-Go StageTips. Purified peptides were analyzed on an Orbitrap Eclipse Tribid Mass Spectrometer (Thermo) coupled with an easy nLC 1200 (Thermo). An in-house packed C18 analytical column (30 cm, 75 μm inside diameter, and 1.9-μm ReproSil-Pur C18 beads; Dr. Maisch) was used with an integrated column oven (50°C; PRSO-V1, Biberach). DiGly-remnant peptides were analyzed using a 90 min gradient (0-72 min: 4-30%B, 73-80 min: 30-55% B, 81-90 min: 55-95% B). Peptides were analyzed in a data-dependent acquisition (DIA) mode. Four full MS scans were set to 120,000 at mass/charge ratio 390 to 1010 m/z, with AGC target at 300% and maximum injection time of 52 ms. DIA scans were performed from a precursor mass range m/z 400-550, 550-700, 700-850, 850-1000 with a resolution of 50,000.

The mass spectrometry proteomics data have been deposited to the ProteomeXchange Consortium via the PRIDE partner repository with the dataset identifier PXD057938^88^. Acquired spectra from DiGly samples were processed using Spectronaut (version 19.1) using library free search against UniProt mouse database (April 2024). Statistical analysis was performed on the MaxLFQ normalized data with R (V 4.2.2), and Perseus (V.1.6.5.0) and visualized with Instant clue (V 0.10.10.20210315). Data completeness of 70% was calculated on each group. Missing values were imputed by random forest algorithm for groups with 70% data completeness, with more than 30% missing values random forest algorithm was downshifted of 0.3, width 1.5. ANOVA analysis and Welch’s T-test analysis was performed with an FDR with less than 0.05 and 500 randomizations.

### RT-qPCR

Total RNA was reverse transcribed into cDNA using the LunaScript RT SuperMix (New England Biolabs, M3010). RT-qPCR was performed using Luna Universal qPCR Master Mix (New England Biolabs, M3003) and run on a LightCycler® 480 System (Roche). Gene expression levels were normalized to the housekeeping gene 18S rRNA. Relative gene expression was determined using a comparative quantification approach and is presented as fold change relative to untreated wt control samples. Primers targeting *Nfκbiα* (Mm.PT.58.21596529), *Tnfaip3* (Mm.PT.58.29149200), *Ccl2* (Mm.PT.58.42151692), *Mmp13* (Mm.PT.58.42286812), *Il6* (Mm.PT.58.10005566) and *Cxcl1* (Mm.PT.58.42076891) were directly purchased from Integrated DNA Technologies, while the primer sequence for 18S rRNA^67^ was obtained from a previous publication.

### Statistics

Data shown represent the mean ± SEM, as indicated in the figure legends. Statistical significance was determined using unpaired, two-tailed parametric Student’s t-test, Log-rank (Mantel-Cox) test, one- or two-way ANOVA with Tukey’s multiple comparisons test with \**p* < 0.05, \*\**p* < 0.01, \*\*\**p* < 0.001, \*\*\*\**p* < 0.0001. All statistical analyses were performed using GraphPad Prism 10, unless stated differently. Statistical details for the proteomic analysis are provided in the preceding section. All in-vitro experiments were performed at least twice with similar results.

---

**Figure S1:**
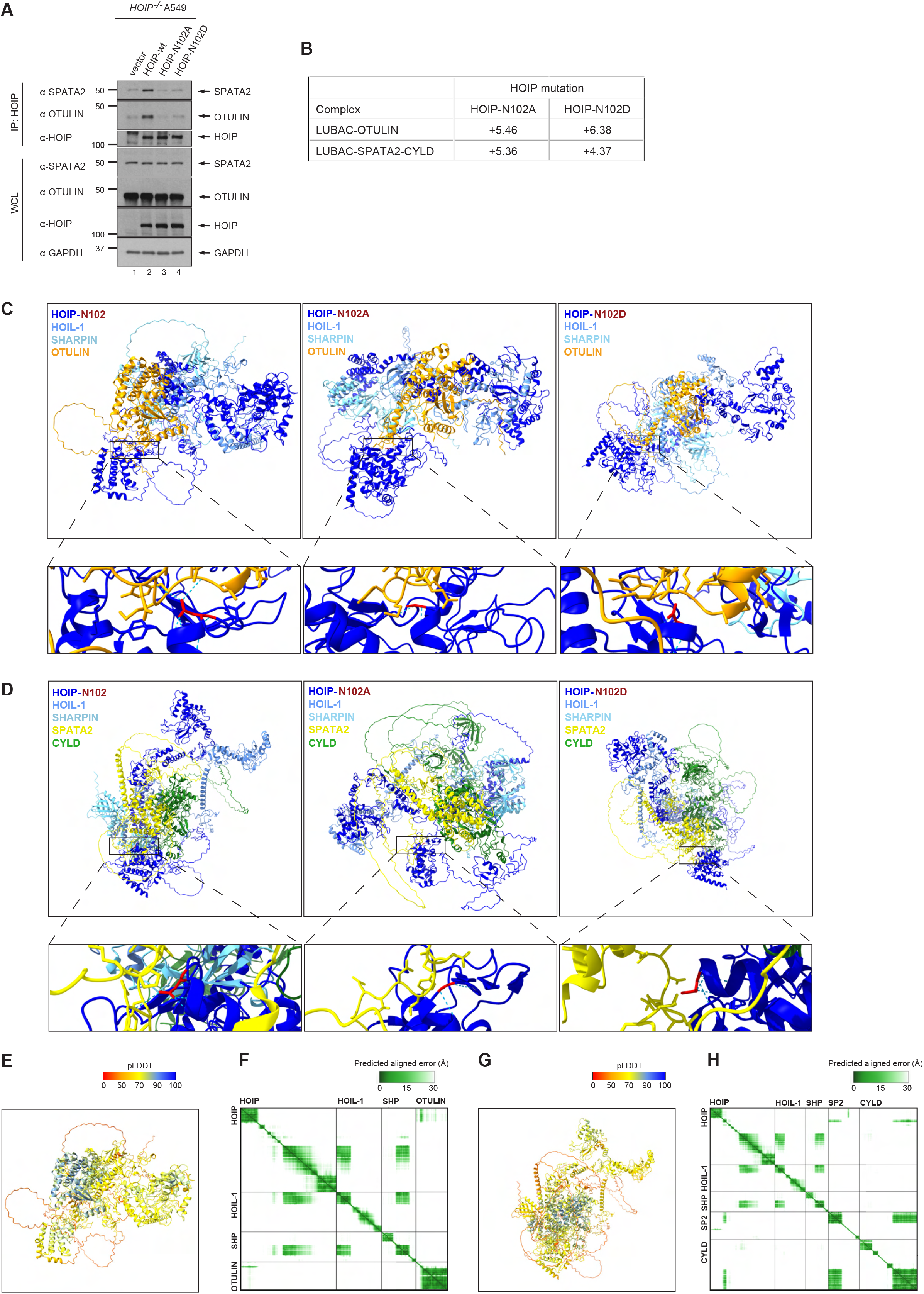
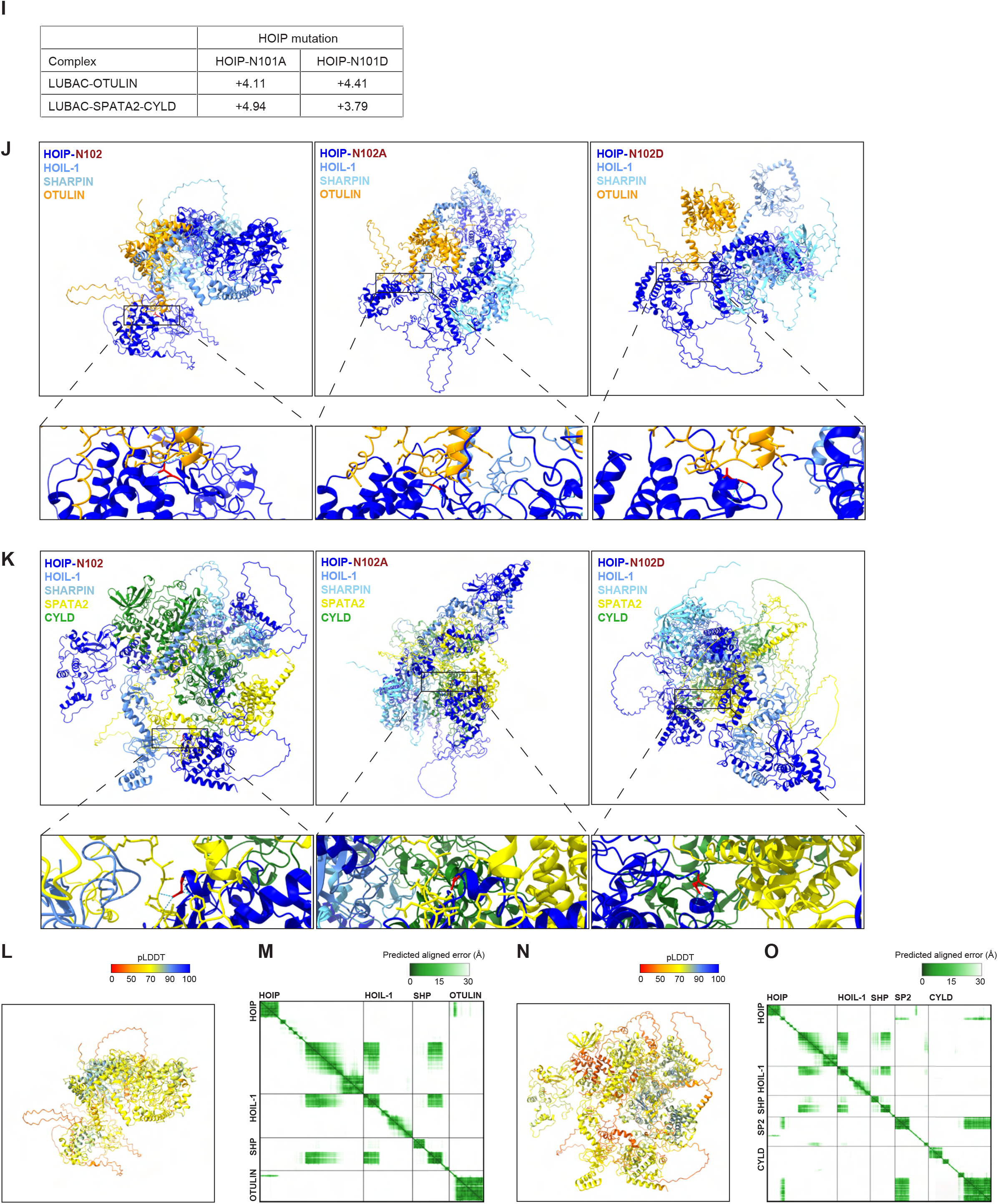
Mutation of HOIP-N102 and HOIP-N101 to alanine or aspartic acid similarly disrupts the HOIP/DUB interaction. (A) *HOIP^-/-^*A549 cells transiently reconstituted with HOIP-wt, HOIP-N102A, HOIP-N102D or empty vector control were immunoprecipitated with anti-HOIP antibody and analyzed by western blot (n=2). (B-O) Structural predictions of the indicated LUBAC components with human HOIP-N102 (B-H) or mouse HOIP-N101 (I-O) variants interacting with OTULIN (C, J) or SPATA2 and CYLD (D, K) via AlphaFold 3. Amino acids proximal to the interaction motif are shown in a stick model. Hydrogen bonds from HOIP-N102 and HOIP-N101 are highlighted as dashed blue lines. FoldX 5.1-predicted ΔΔG (kcal/mol) for HOIP mutations were calculated with BuildModel after RepairPDB preprocessing (B, I). Positive values indicate destabilization relative to wild-type HOIP upon mutation to N102A or N102D in the indicated complexes. Representative predicted structures of the wt LUBAC–DUB complexes are shown colored by pLDDT confidence scores (E, G, L, N) and the predicted aligned error (PAE) plots of the corresponding wt complexes are shown in (F, H, M, O).

**Figure S2:**
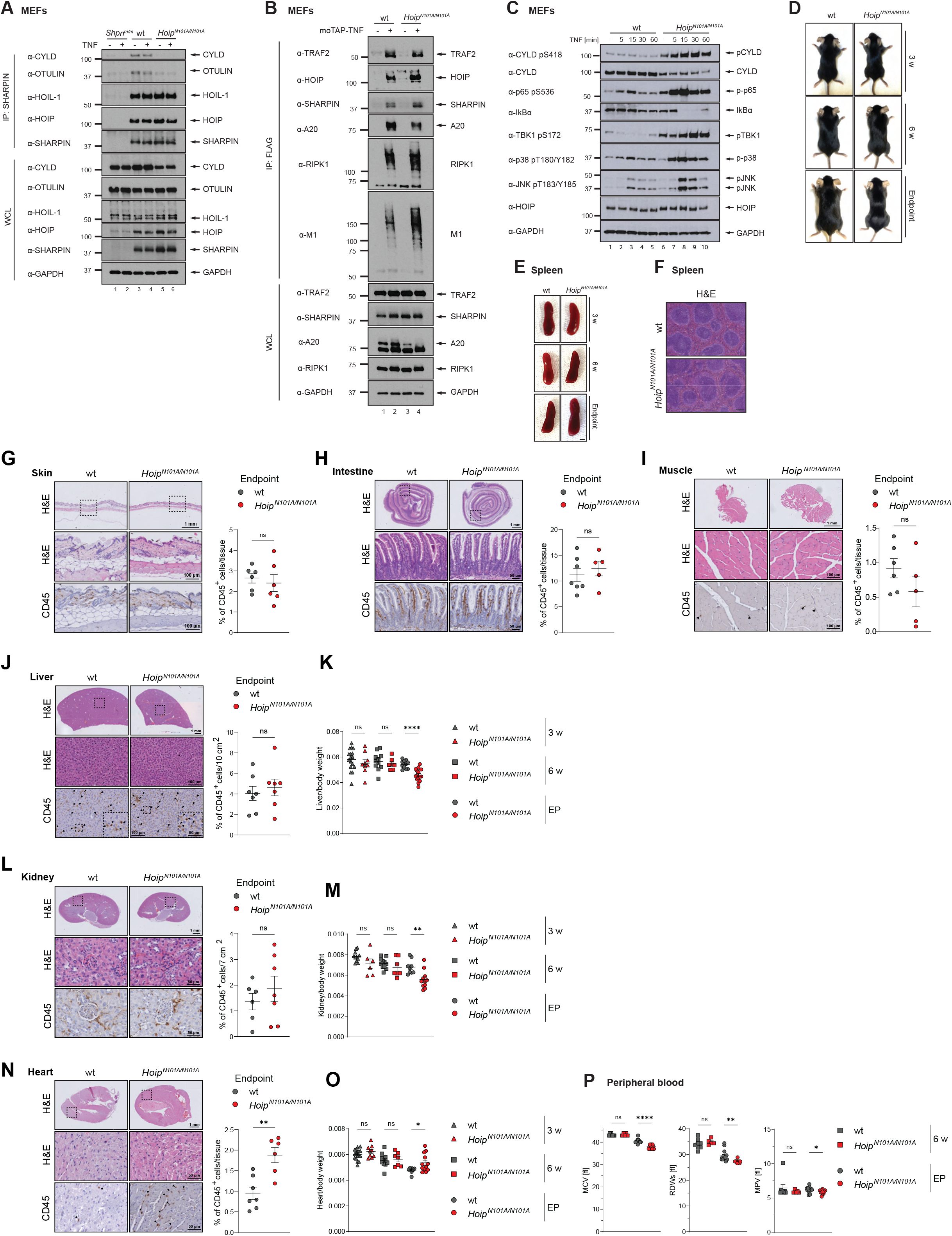
*N101A* mice show mild cardiac inflammation and reduced blood oxygenation. (A) MEFs were stimulated with 100 ng/ml TNF for 5 min or left untreated. Lysates were subjected to immunoprecipitation using an anti-SHARPIN antibody and analyzed via western blot (n=2). (B) MEFs were treated with 1 μg/ml moTAP-TNF for 5 min or left untreated. Lysates were immunoprecipitated with anti-FLAG sepharose beads and analyzed via western blot (n=2). (C) Western blot analysis of MEFs stimulated with 100 ng/ml TNF for the indicated time points (n=2). (D, E) Representative images of male mice (E) and spleens (E) with indicated genotype and age. Scale bars: 200 μm or 1 cm respectively. (F) Representative H&E staining of spleen (n=5/genotype). Scale bar: 100 μm. (G-O) H&E, CD45 IHC staining and organ-to-body-weight ratios from *N101A* mice sacrificed at humane endpoint and age-matched wt controls. (P) Peripheral blood analysis performed with differential hematology analyzer at six weeks of age and survival endpoint. Values are shown for the mean corpuscular volume (MCV), red cell distribution width standard deviation (RDWs) and mean platelet volume (MPV). Data present mean ± SEM. Statistical analyses were performed via two-tailed unpaired t-test with \**p* < 0.05, \*\**p* < 0.01, \*\*\**p* < 0.001, \*\*\*\**p* < 0.0001.

**Figure S3:**
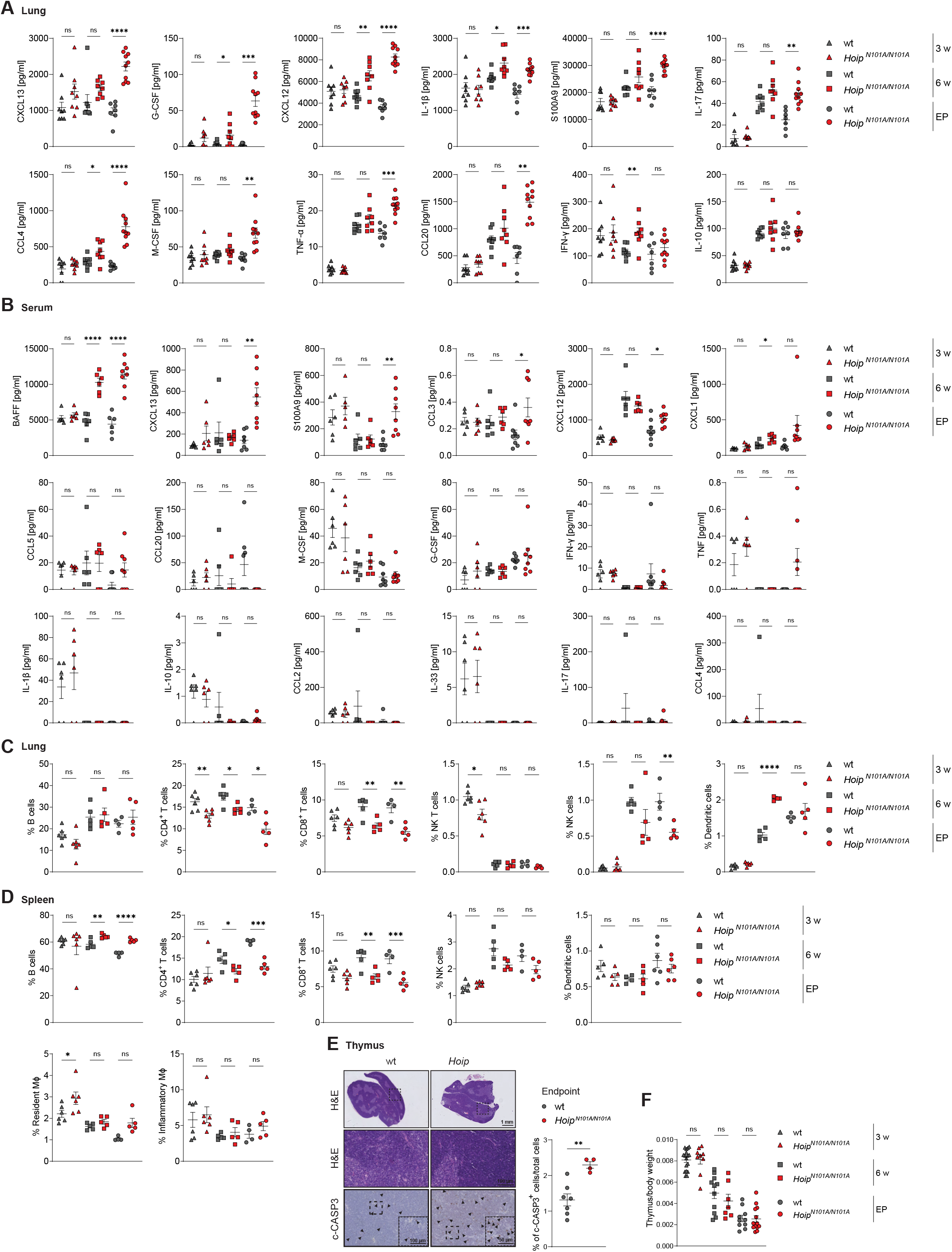
HOIP-N101A induces progressive, severe inflammatory lung pathology. (A, B) Cytokine levels of lung homogenates (A) and serum (B) were analyzed using a Luminex-Multiplex assay for the indicated targets. (C, D) Flow cytometric analysis of lungs (C) and spleens (D) at indicated ages. All immune cell subsets represent the percentage from viable CD45^+^ cells. MΦ = macrophages. (E) H&E and IHC staining for cleaved Caspase-3 (c-CASP3) of thymus from *N101A* mice sacrificed at the humane endpoint or age-matched wt controls. (F) Thymus-to-body weight ratio of indicated genotypes and age. All data present mean ± SEM. Statistical analyses were performed via two-tailed unpaired t-test with \**p* < 0.05, \*\**p* < 0.01, \*\*\**p* < 0.001, \*\*\*\**p* < 0.0001.

**Figure S4:**
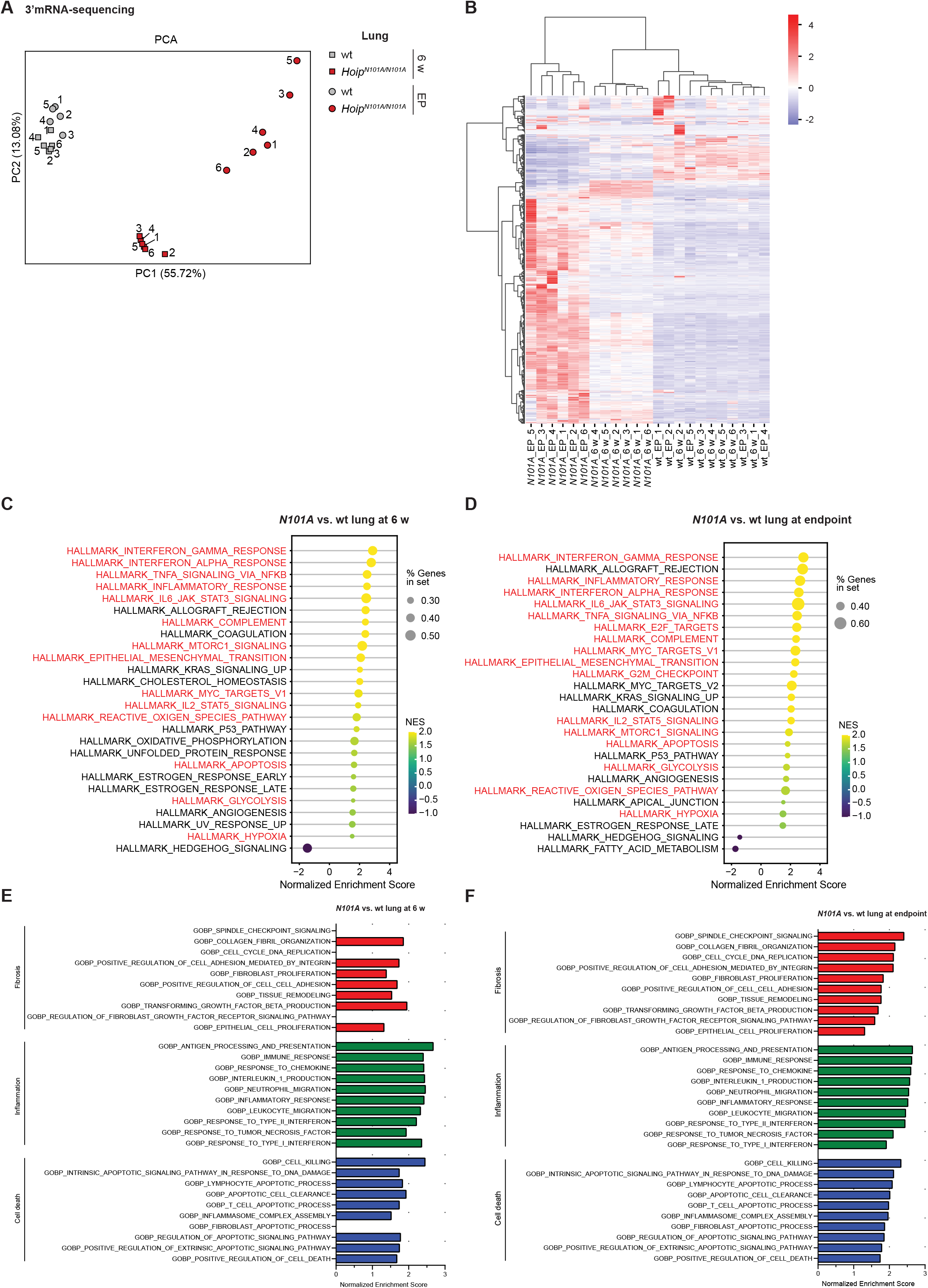
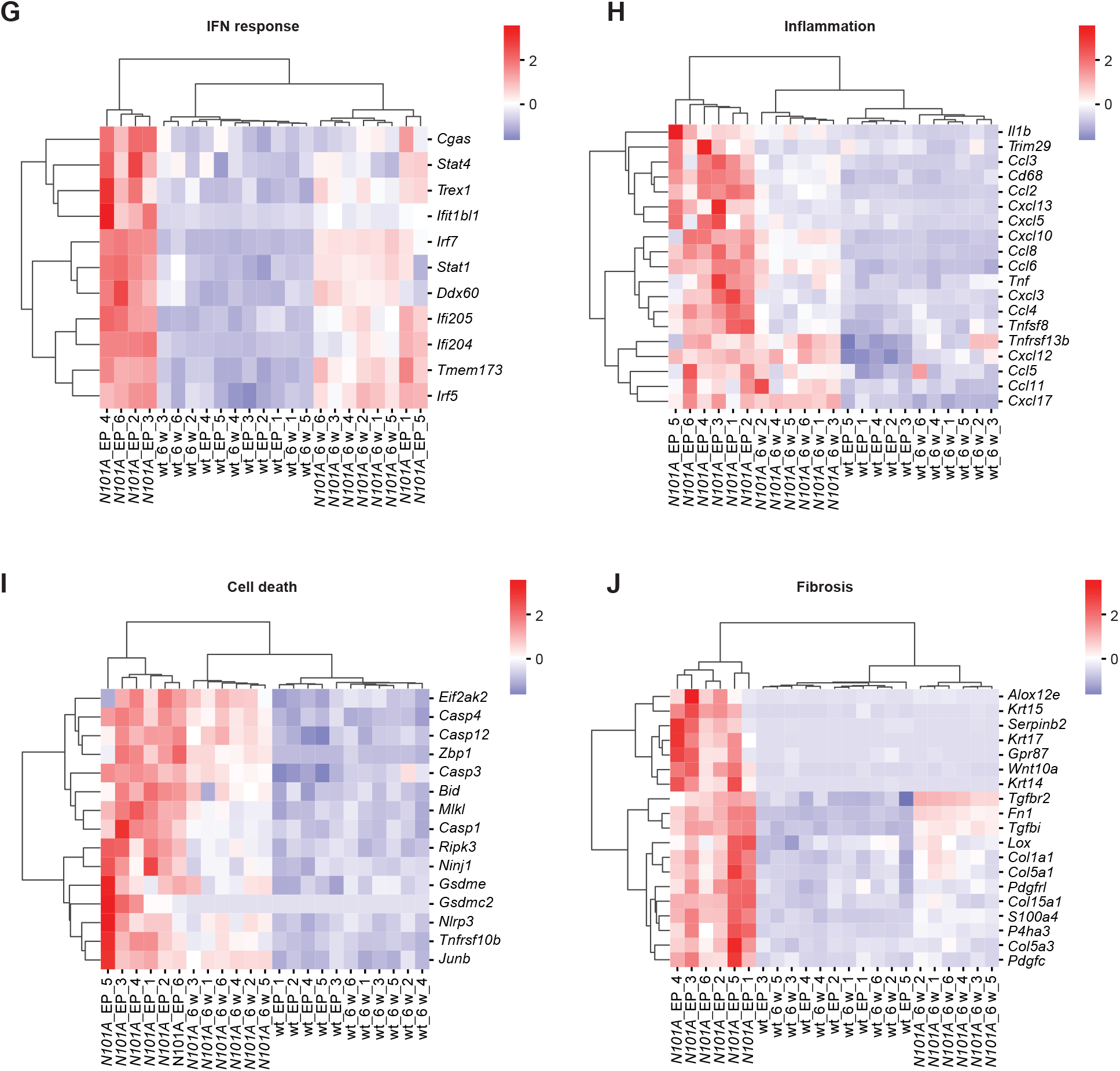
HOIP-N101A exacerbates gene transcription promoting pro-fibrotic, pro-inflammatory and cell-death–associated gene expression. (A) Lungs of the indicated genotypes and time points were subjected to 3’ mRNA-sequencing. The principal component analysis (PCA) illustrates the separation of samples based on transcriptomic profiles. (B) Cluster heatmap of significantly differentially expressed genes across different genotypes and time points. (C, D) Unbiased hallmark analysis of significantly altered pathways of *N101A* lung samples normalized on wt controls at six-weeks of age (C) or survival endpoint (D). (E, F) Pathway enrichment analysis based on Gene Ontology Biological Process (GO BP) terms. Selected significantly upregulated pathways of *N101A* samples normalized on wt controls at six-weeks of age (E) or survival endpoint (F). (G-J) Cluster heatmaps show significantly differentially expressed genes based on the statistical difference between *N101A* and wt lungs at survival endpoint. The color intensity represents log2Fold changes in gene expression.

**Figure S5:**
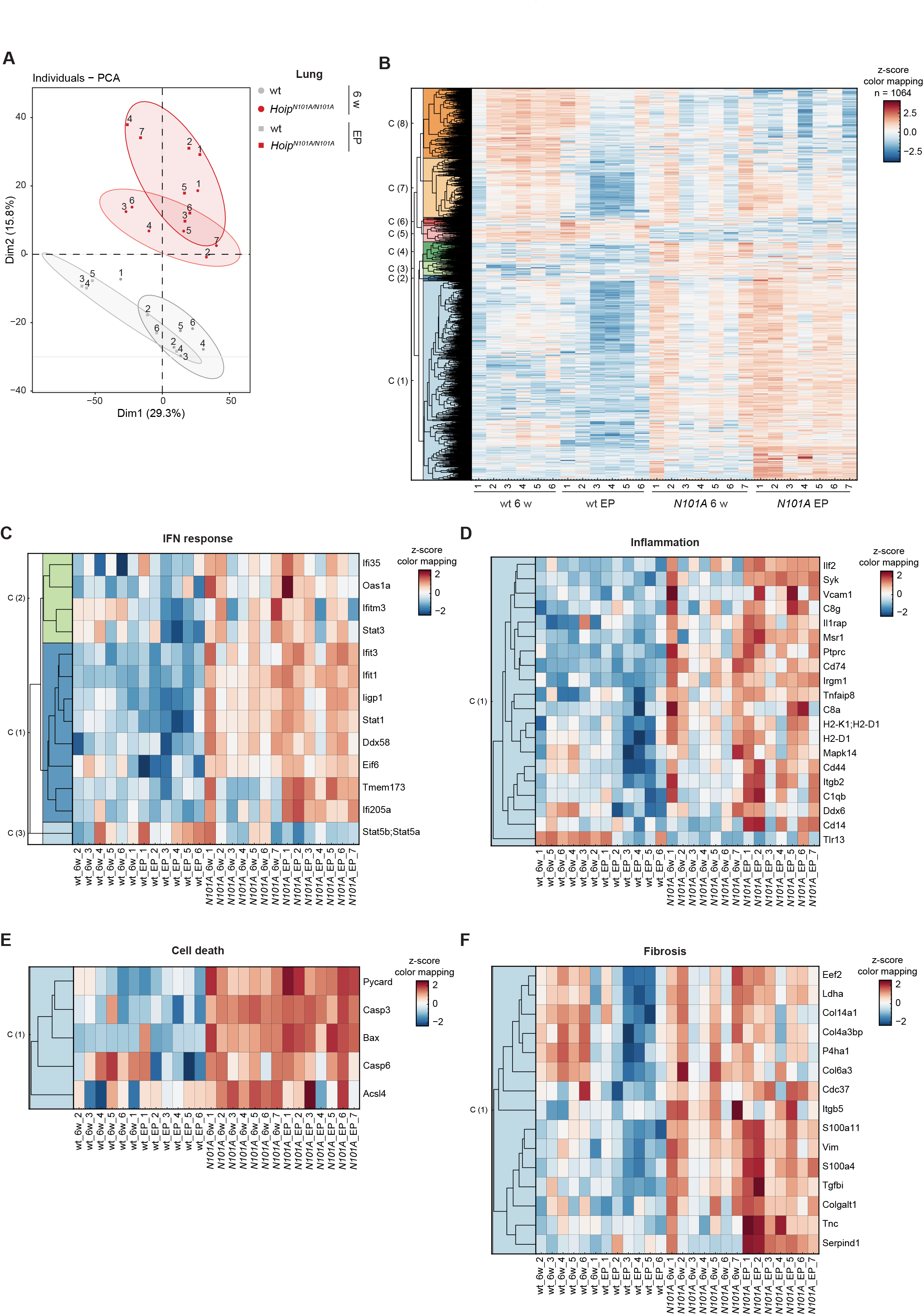
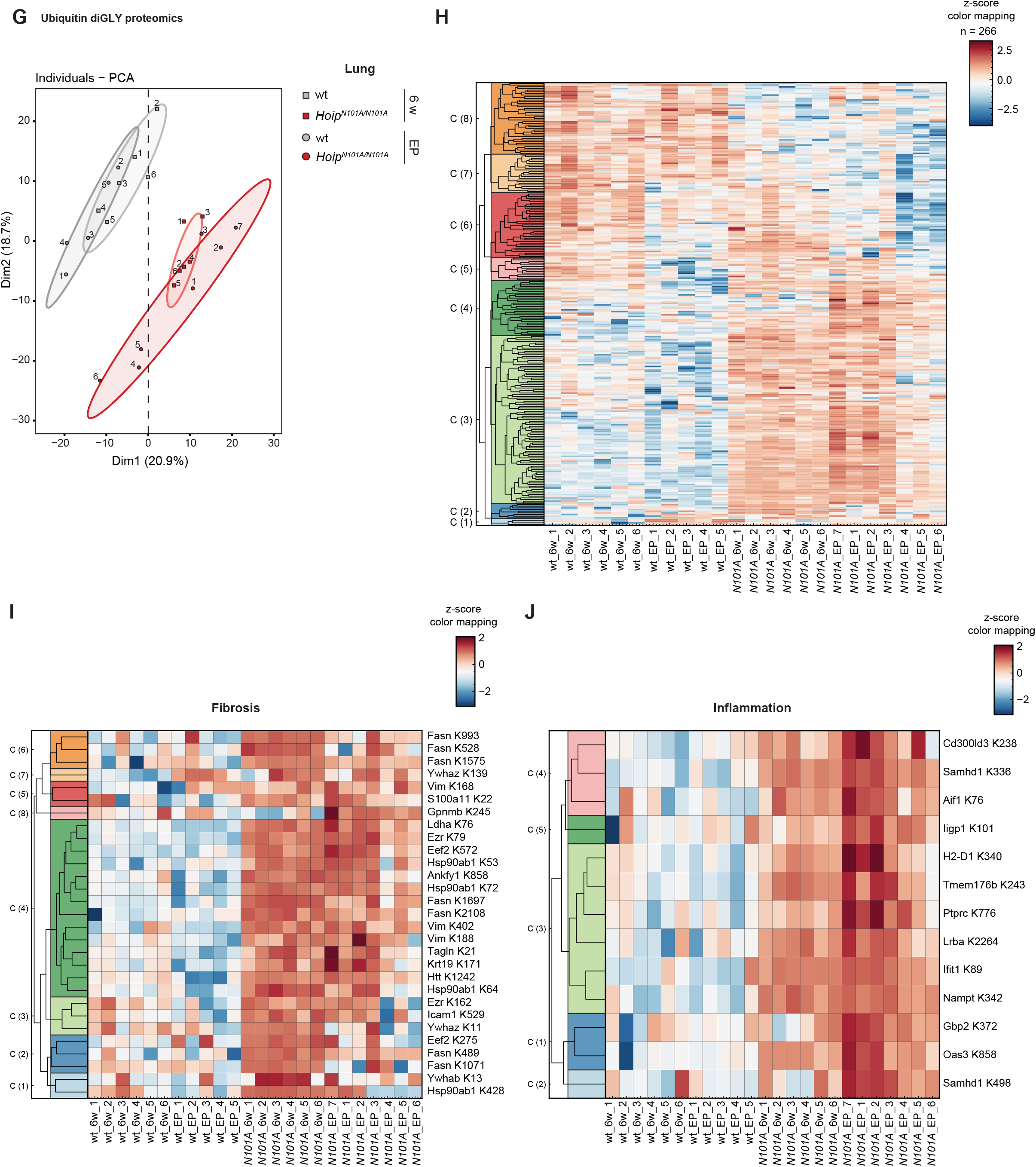
Proteomic profiling of *N101A* lungs reveals increased pro-fibrotic proteins and ubiquitination of pro-fibrotic and pro-inflammatory proteins. (A) Lungs of the indicated genotypes and time points were subjected to proteomic analysis. Individuals-Principal component analysis (PCA) illustrating the separation of samples based on proteomic profiles. (B-F) Cluster heatmaps showing the significantly different abundance of proteins across indicated genotypes and time points. The color intensity indicates changes in protein enrichment expressed as z-scores. Statistical analysis was performed via one-way ANOVA. (G-J) Ubiquitin diGly proteomics were performed with the same lung homogenates. Principal component analysis (PCA) of all differentially ubiquitinated proteins is presented (G) and cluster heatmaps of significantly differentially ubiquitinated proteins across genotypes and time points are shown (H-J). The color intensity indicates the level of di-glycine remnant enrichment as z-scores. Statistical analysis was performed via one-way ANOVA.

**Figure S6:**
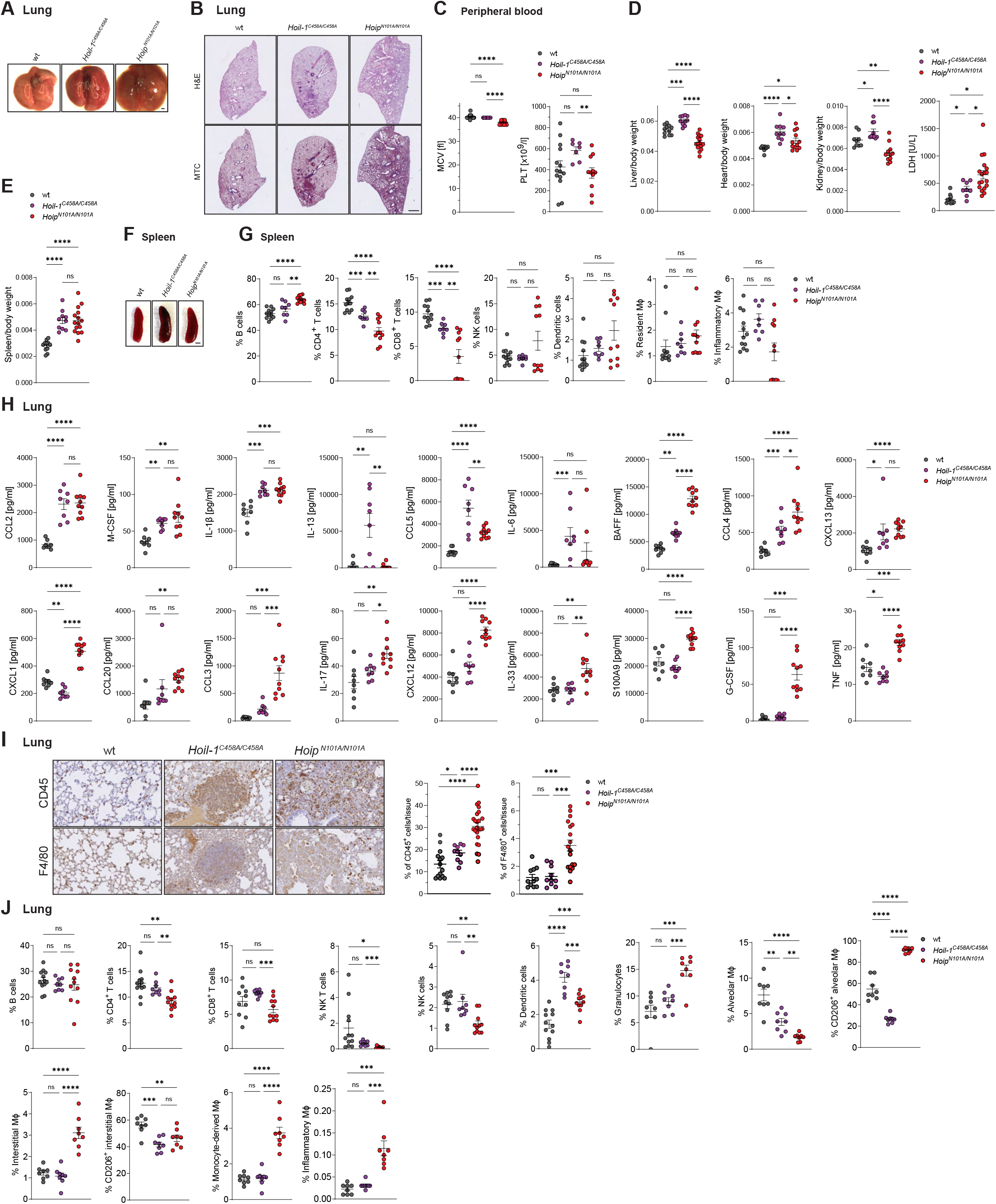
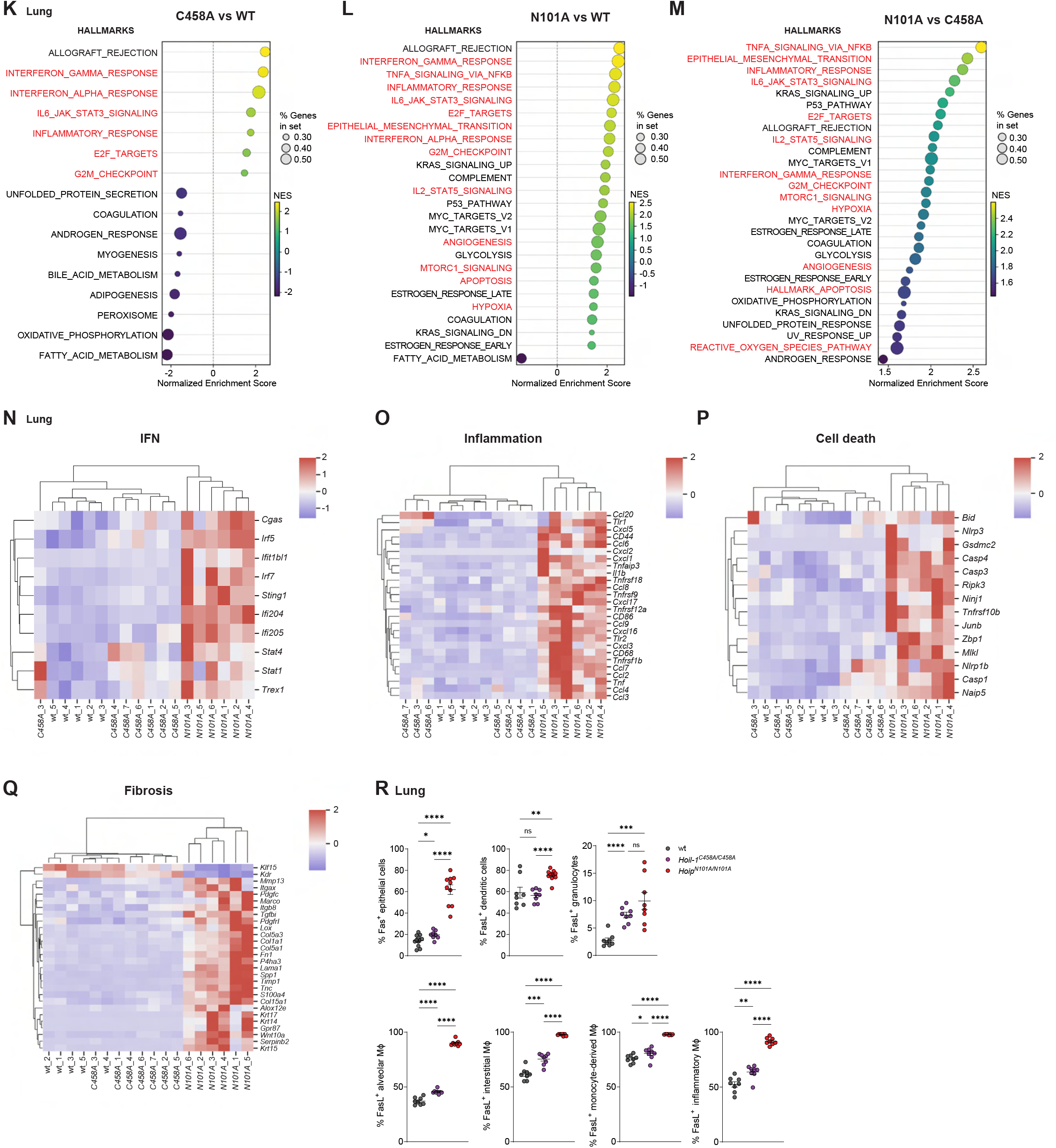
*C458A* mice exhibit mild systemic inflammation without overt pathology. (A, B) Body weight and spleen-to-body weight ratio with representative images of spleens. Scale bar: 200 µm. (C) Flow cytometric analysis of spleen from mice of indicated genotypes at 10 weeks of age. MΦ = macrophages. Data present mean percentages of viable CD45^+^ cells ± SEM (n ≥ 8 mice per genotype). (D) Organ-to-body weight ratios and serum LDH levels. (E, F) Representative images of lungs (E) and overview images of H&E and MTC stainings of lung sections (F). Scale bars: 1 cm and 100 μm respectively. (E) Mean corpuscular volume (MCV) and platelets (PLT) in the peripheral blood. (H) Cytokine levels of lung homogenates were analyzed using a Luminex-Multiplex assay for the indicated targets. (I) IHC staining of lungs for CD45 and F4/80 with corresponding quantifications. Scale bar: 100 μm. (J) Flow cytometric analysis of lungs from mice of indicated genotypes at 10 weeks of age. Data present mean percentages of viable CD45^+^ cells (n ≥ 8 mice per genotype). MΦ = macrophages. (K-M) Unbiased hallmark analysis of significantly altered pathways of *C458A*, wt and *N101A* lung samples normalized as indicated. (N-Q) Cluster heatmaps show significantly differentially expressed genes based on the statistical difference between *N101A* and wt lungs. The color intensity represents log2Fold changes in gene expression. (R) Flow cytometric analysis of lungs from mice of indicated genotypes at 10 weeks of age. The percentages of Fas^+^ and FasL^+^ cells are shown as the proportion of positive cells within the parental population. MΦ = macrophages. Data present mean ± SEM. Statistical analyses were performed via two-tailed unpaired t-test with \**p* < 0.05, \*\**p* < 0.01, \*\*\**p* < 0.001, \*\*\*\**p* < 0.0001. If not otherwise indicated, wt and *C458A* mice were sacrificed at ∼15 weeks of age, *N101A* mice at their respective humane endpoint (median survival ∼15 weeks).

**Figure S7:**
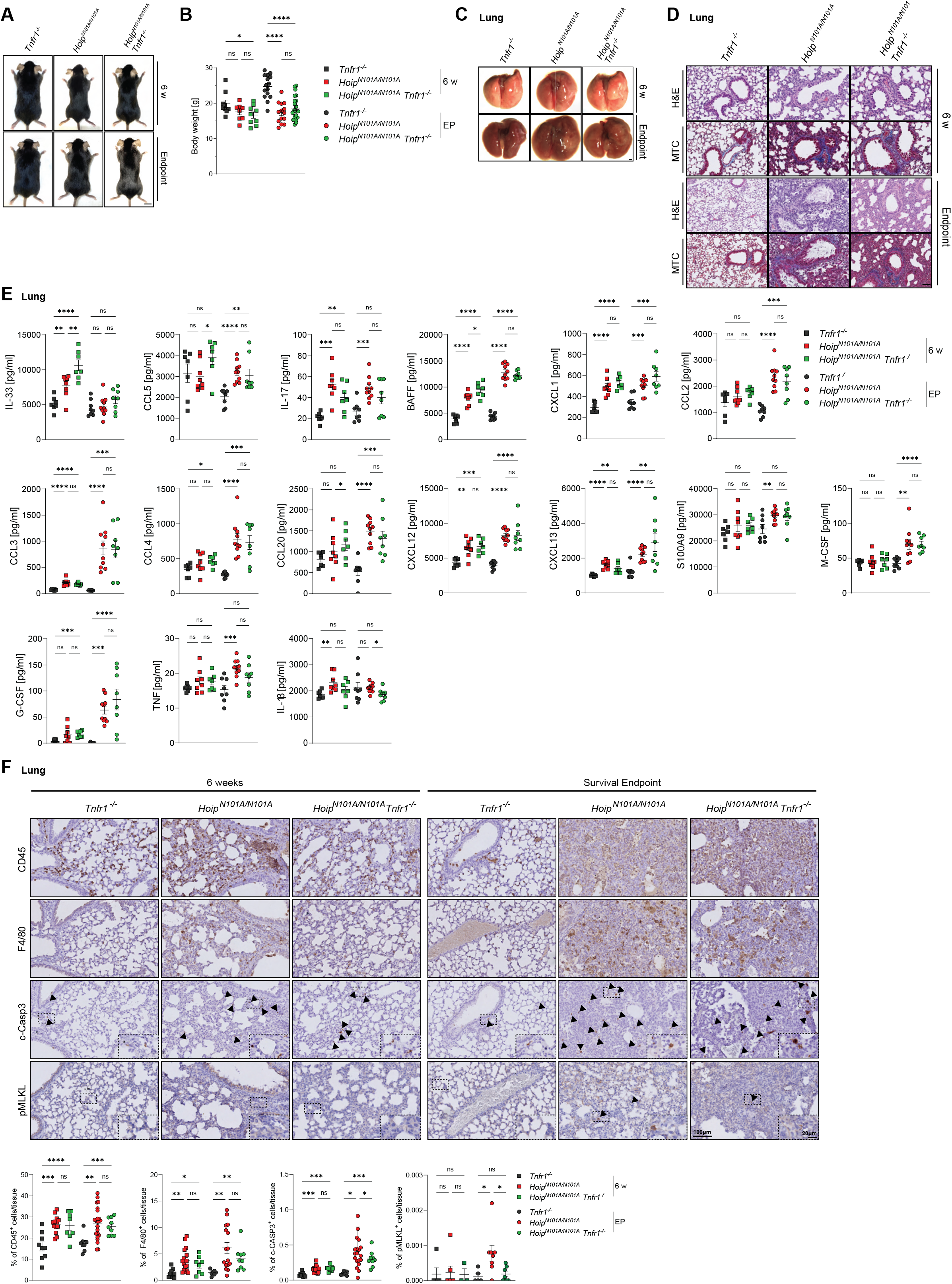
TNFR1 contributes to cell death in *N101A* lungs. (A, B) Representative images of male mice (A) and body weight (B) of indicated genotypes and ages. Scale bar: 1 cm. (C) Representative images of lungs. Scale bar: 1 cm. (D) H&E and MTC staining of lung. Scale bar: 100 μm. (E) Cytokine levels of lung homogenates were analyzed using a Luminex-Multiplex assay for the indicated targets. (F) IHC stainings of lungs for CD45, F4/80, cleaved Caspase-3 (c-CASP3) and phospho-MLKL (MLKL pS345) with the respective quantifications. Scale bar: 100 μm. Data present mean ± SEM. Statistical analyses were performed via two-tailed unpaired t-test with \**p* < 0.05, \*\**p* < 0.01, \*\*\**p* < 0.001, \*\*\*\**p* < 0.0001.

**Figure S8:**
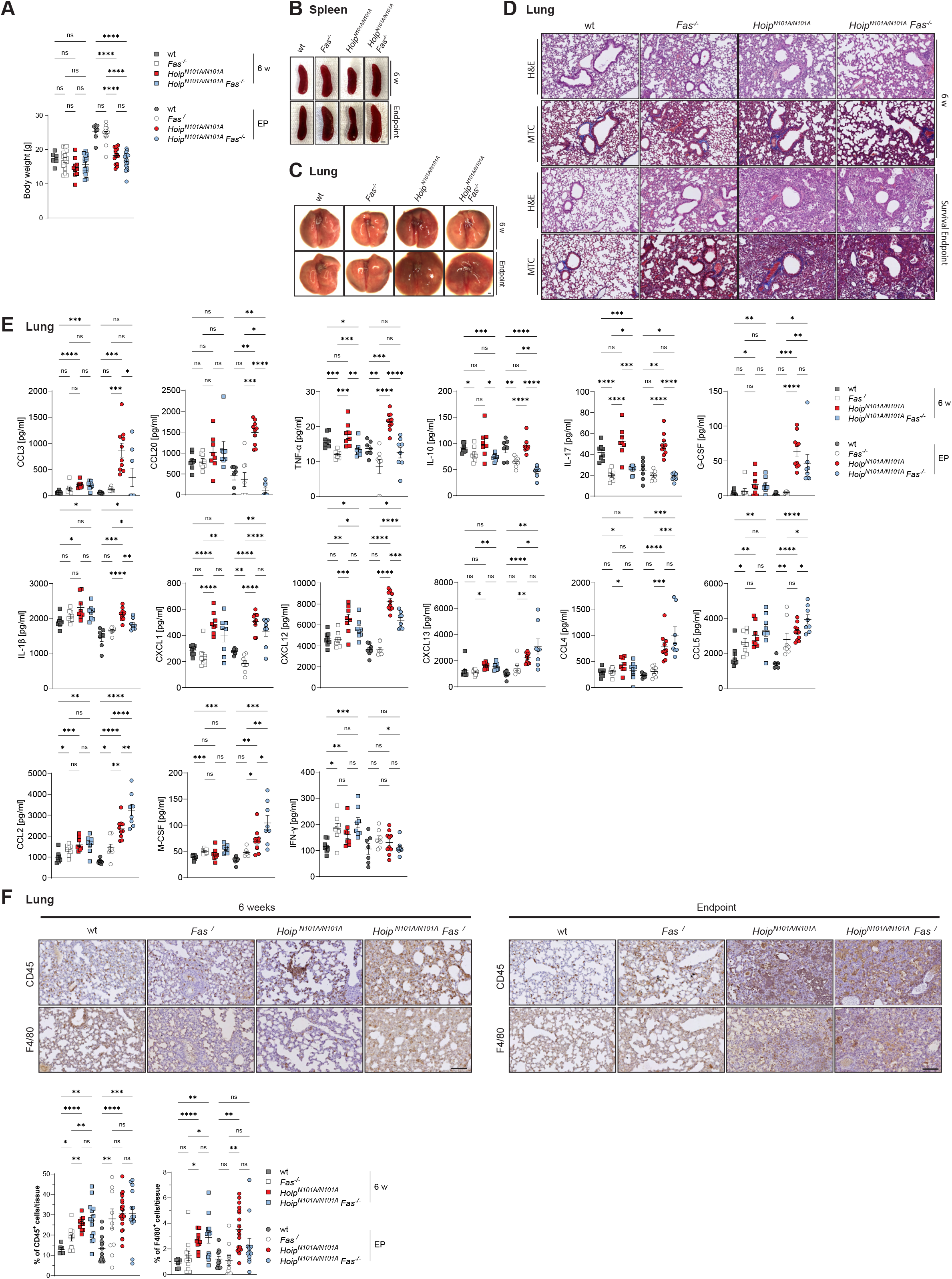
Fas contributes to the inflammatory milieu in *N101A* lungs. (A) Body weight of indicated genotypes and ages. (B, C) Representative images of spleens (B) and lungs (C). Scale bar: 200 μm and 1 cm. (D) H&E and MTC staining of lung. Scale bar: 100 μm. (E) Cytokine levels of lung homogenates were analyzed using a Luminex-Multiplex assay for the indicated targets. (F) IHC stainings of lungs for CD45 and F4/80 with the respective quantifications. Scale bar: 100 μm. Data present mean ± SEM. Statistical analyses were performed via two-tailed unpaired t-test with \**p* < 0.05, \*\**p* < 0.01, \*\*\**p* < 0.001, \*\*\*\**p* < 0.0001.

**Figure S9:**
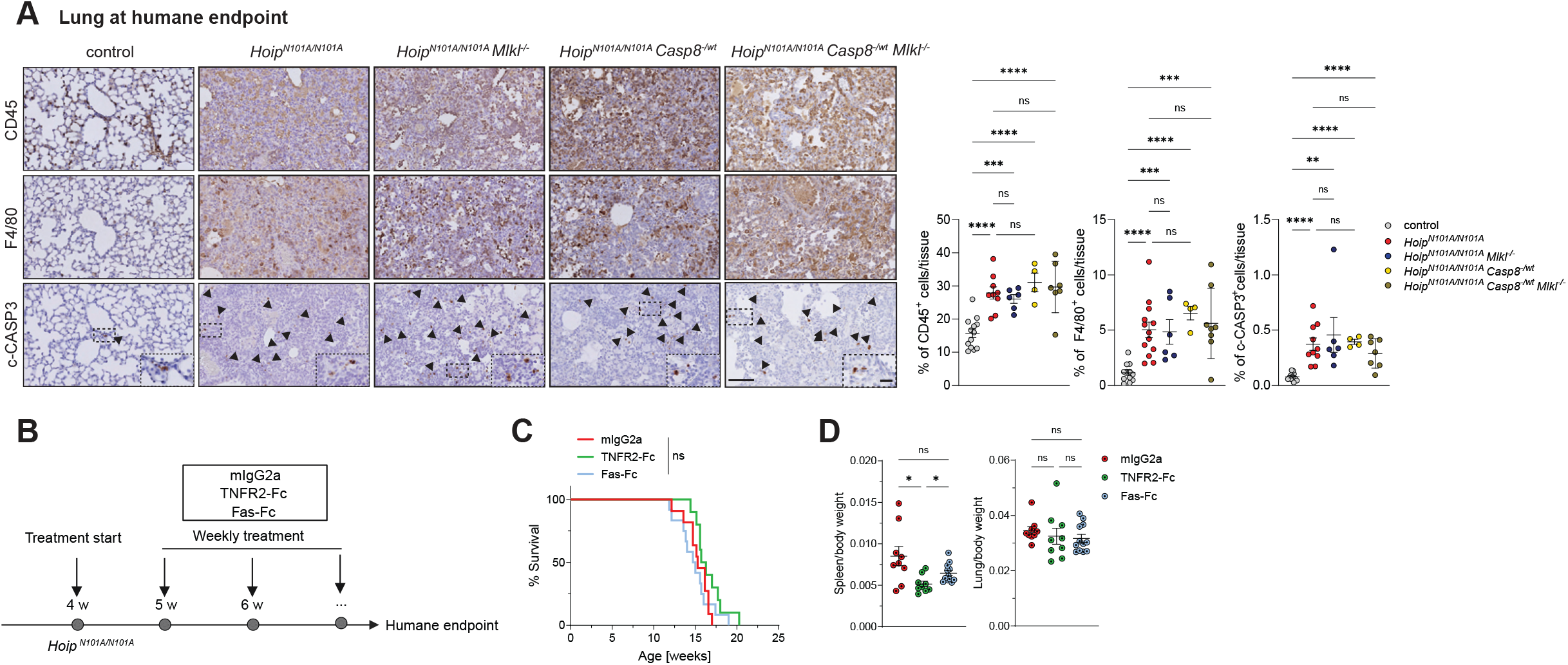
Partial cell death suppression does not prevent lung pathology in *N101A* mice. (A) IHC stainings of lungs for CD45, F4/80 and cleaved Caspase-3 (c-CASP3) with the respective quantifications. Scale bar: 100 μm. (B) Experimental design: 4-week-old *N101A* mice were randomized into the indicated treatment groups. 500 µg of each compound was administered intraperitoneally once per week from 4 weeks of age until the humane endpoint. (C) Kaplan-Meier survival analysis of *N101A* mice treated with the indicated Fc proteins. Statistical analysis was performed via Log-rank (Mantel-Cox) test with \**p* < 0.05. (D) Organ-to-body-weight ratios. All data present mean ± SEM. Statistical analysis was performed via two-tailed unpaired t-test with \**p* < 0.05, \*\**p* < 0.01, \*\*\**p* < 0.001, \*\*\*\**p* < 0.0001.

**Figure S10:**
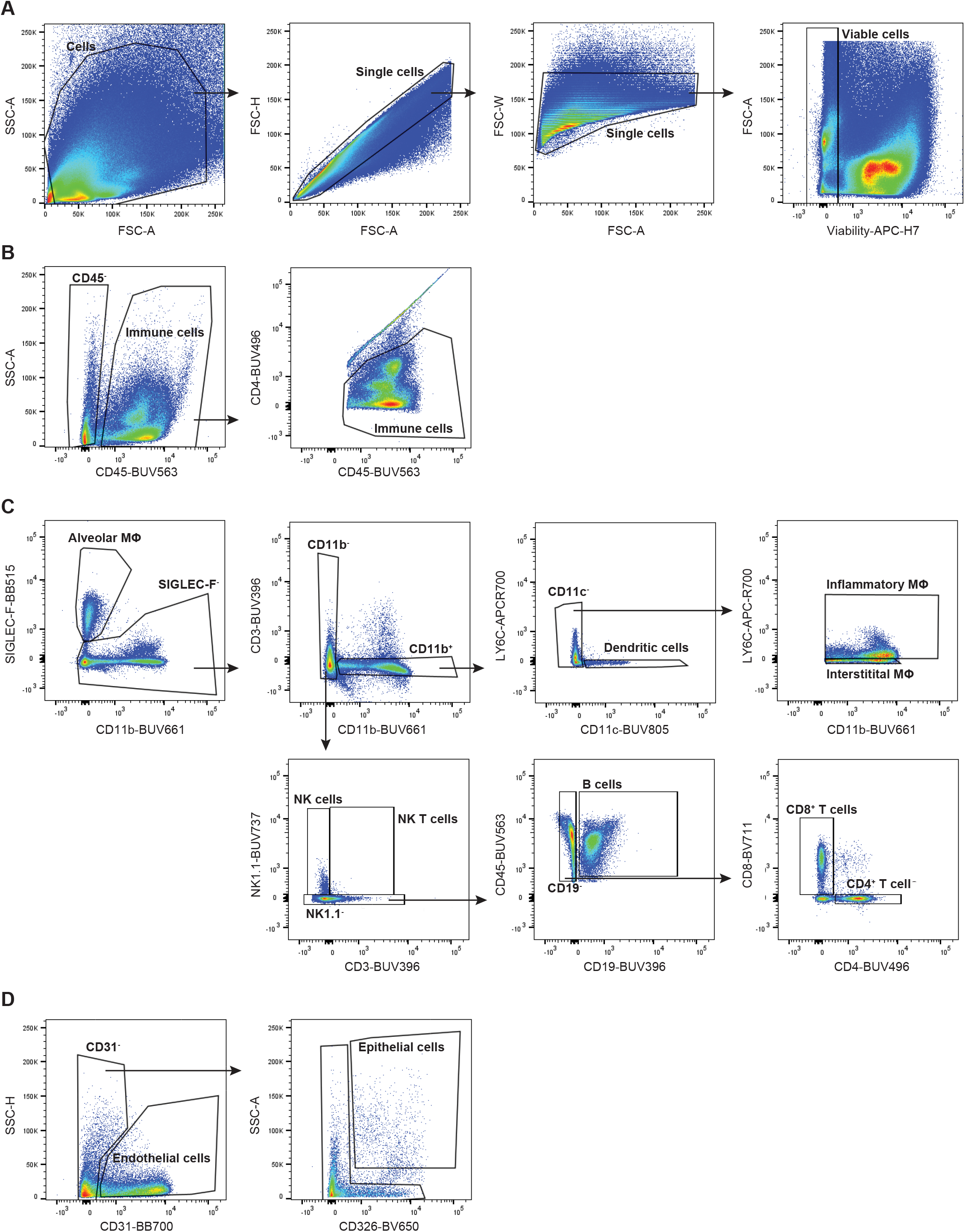
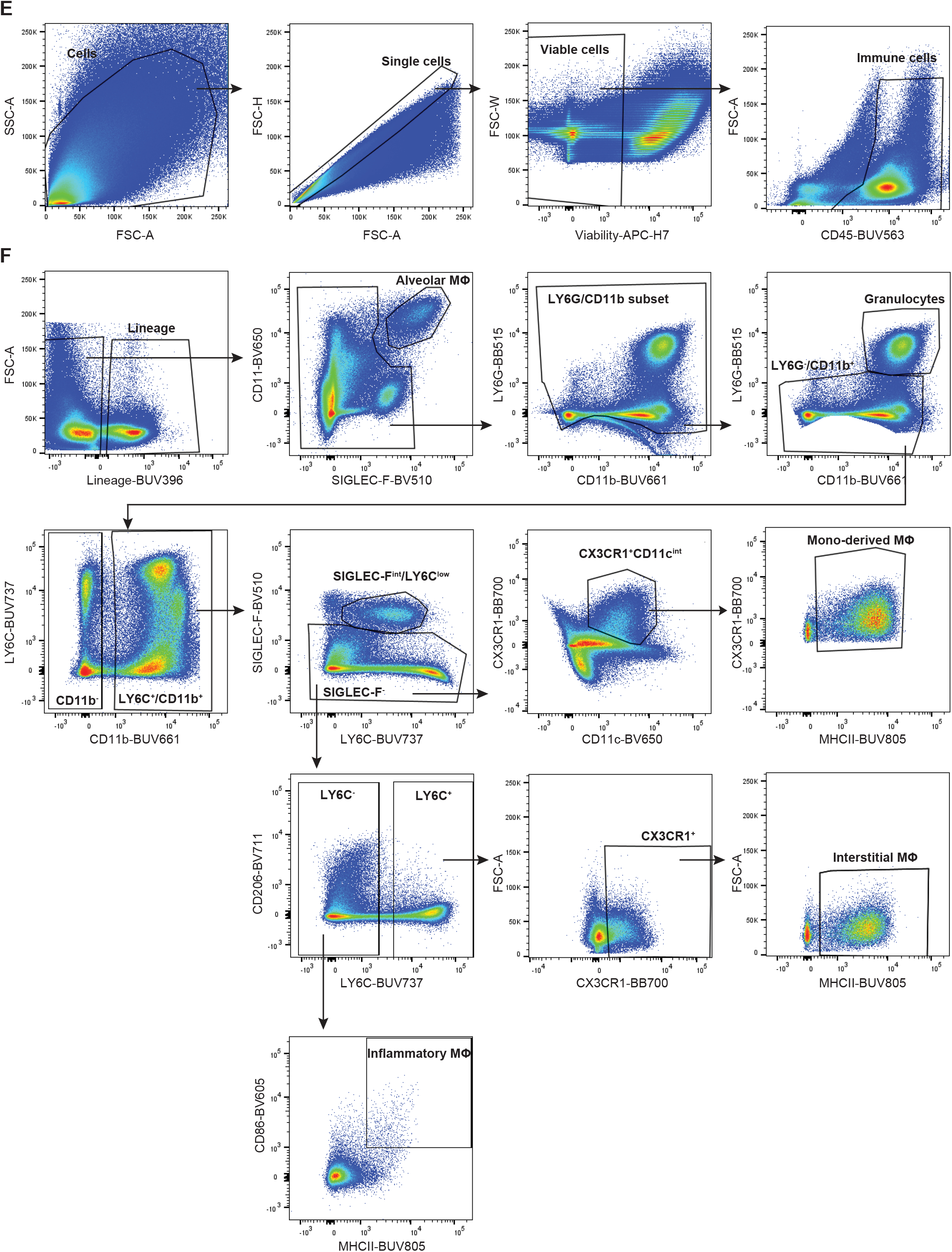
Gating strategy for identification of immune and non-immune cells from mouse lung and spleen. (A-D) Gating strategy for identification of immune and non-immune cells from mouse lung and spleen. (A) Selection of viable single cells. (B) Identification of immune cells (CD45⁺). (C) Gating of distinct immune cell subsets within the CD45⁺ population. (D) Identification of non-immune cell populations within the CD45⁻ fraction. (E-F) Gating strategy for identification of distinct myeloid cell populations from mouse lung. (E) Selection of viable single immune cells. (F) Gating of distinct myeloid cell subsets within the CD45⁺ population.

## SUPPLEMENTARY MATERIALS

**Table S1.** Primary antibodies used for immunoblotting.

| Antibodies | Source | Identifier |
| --- | --- | --- |
| A20 | Cell Signaling Technology | Cat# 5630T |
| α-TUBULIN | Bio-Rad | Cat# MCA78G |
| c-CASP3 (D175) | Cell Signaling Technology | Cat# 9664 |
| CYLD | Santa Cruz Biotechnology | Cat# sc-74435 |
| GAPDH | Sigma-Aldrich | Cat# G9545 |
| HOIL-1 | self-produced |  |
| HOIP | abcam | Cat# 46322 |
| IκBα | Cell Signaling Technology | Cat# 4812 |
| linUb | Sigma-Aldrich | Cat# ZRB2114 |
| OTULIN | abcam | Cat# ab211328 |
| Phospho-CYLD (S418) | Cell Signaling Technology | Cat# 4500 |
| Phospho-ERK1/2 (T202/Y204) | Cell Signaling Technology | Cat# 4370 |
| Phospho-JNK (T183/Y185) | Cell Signaling Technology | Cat# 4668S |
| Phospho-MLKL (S345) | Cell Signaling Technology | Cat# 37333 |
| Phospho-p38 (T180/Y182) | Cell Signaling Technology | Cat# 4511 |
| Phospho-p65 (S536) | Cell Signaling Technology | Cat# 3033 |
| Phospho-TBK1/NAK (S172) | Cell Signaling Technology | Cat# 5483 |
| RIP1 | Cell Signaling Technology | Cat# 3493 |
| RIP3 | Cell Signaling Technology | Cat# 95702 |
| SHARPIN | Proteintech | Cat# 14626-I-AP |
| SPATA-2 | Bethyl Laboratories | Cat# A302-494A |
| TNFR1 | abcam | Cat# ab19139 |
| TRAF-2 | Santa Cruz Biotechnology | Cat# sc-876 |

**Table S2.** Antibodies for flow cytometric analysis of lungs and spleens.

| **Antibody** | **Source** | **Identifier** | **Dilution** |
| --- | --- | --- | --- |
| CD11b | BD Biosciences | Cat# 612977 | 1:250 |
| Cd11c | BioLegend | Cat# 117334 | 1:100 |
| CD19 | BD Biosciences | Cat# 565965 | 1:200 |
| CD206 | BD Biosciences | Cat# 141727 | 1:100 |
| CD3 | BioLegend | Cat# 100234 | 1:200 |
| CD31 | BD Biosciences | Cat# 558738 | 1:100 |
| CD326 | BD Biosciences | Cat# 740559 | 1:100 |
| CD4 | BD Biosciences | Cat# 612952 | 1:250 |
| CD45 | BD Biosciences | Cat# 612924 | 1:250 |
| CD8 | BD Biosciences | Cat# 563046 | 1:250 |
| CD95 | BD Biosciences | Cat# 557653 | 1:100 |
| CD95L | BioLegend | Cat# 106603 | 1:100 |
| CX3CR1 | BD Biosciences | Cat# 567821 | 1:100 |
| Ly6C | BioLegend | Cat# 128041 | 1:150 |
| Ly6G | BD Biosciences | Cat# 741994 | 1:100 |
| MHCII | BD Biosciences | Cat# 748844 | 1:100 |
| NK1.1 | BD Biosciences | Cat# 741715 | 1:250 |

